# Synergistic combination therapy with ONC201 or ONC206, Enzalutamide and Darolutamide in preclinical studies of castration-resistant prostate cancer

**DOI:** 10.1101/2024.07.31.606054

**Authors:** Laura Wu, Maximilian Pinho-Schwermann, Lanlan Zhou, Leiqing Zhang, Kelsey E. Huntington, Ryan Malpass, Attila A. Seyhan, Benedito A. Carneiro, Wafik S. El-Deiry

**Affiliations:** Laboratory of Translational Oncology and Experimental Cancer Therapeutics, The Warren Alpert Medical School, Brown University, Providence, RI, 02903, USA; Biotechnology Graduate Program, Brown University, Providence, RI, 02903, USA; The Joint Program in Cancer Biology, Brown University and the Lifespan Health System, Providence, RI 02903, USA; Department of Pathology and Laboratory Medicine, The Warren Alpert Medical School, Brown University, Providence, RI, 02903, USA; Pathobiology Graduate Program, The Warren Alpert Medical School, Brown University, Providence, RI, 02903, USA; Hematology-Oncology Division, Department of Medicine, Rhode Island Hospital and Brown University, Providence, RI 02903, USA; Legorreta Cancer Center at Brown University, The Warren Alpert Medical School, Brown University, Providence, RI, 02903, USA

**Keywords:** Prostate cancer, ONC201, ONC206, enzalutamide, darolutamide, Castrate resistance

## Abstract

Androgen receptor (AR) signaling plays a primary role in prostate cancer progression. Non-steroidal anti-androgens (NSAA) including enzalutamide, and apalutamide have been used to treat patients with advanced disease. However, patients with metastatic castration-resistant prostate cancer (mCPRC) develop resistance, resulting in limited overall survival benefit. Darolutamide is a novel next-generation androgen receptor-signaling inhibitor that is FDA approved for non-metastatic castration resistant prostate cancer (nmCRPC). Imipridone ONC201/TIC10 is first-in-class small molecule that activates the integrated stress response (ISR) and upregulates TNF-related apoptosis-inducing ligand (TRAIL). Our study investigates ISR and AR signaling in anti-tumor efficacy with ONC201 and enzalutamide or darolutamide against mCRPC cells. mCRPC cell lines 22RV1, LNCaP, DU145 and PC3 were treated with ONC201, darolutamide, and enzalutamide as single agents or in combinations. Combinations of ONC201 and darolutamide or enzalutamide demonstrated synergistic effects in mCRPC cells. Combinations of ONC201 and darolutamide or enzalutamide reduced PSA levels in LNCaP cells and induced of ATF4 in both LNCaP and 22RV1 cell lines. Darolutamide synergized with ONC201 regardless of AR status or castration sensitivity *in vitro.* Flow cytometric analysis showed increased intra-tumoral NK cells in mice treated with ONC201 and combination of ONC201 and darolutamide. Trends of increased TRAIL activation within NK cells were also observed in treatment groups. ONC201 and darolutamide demonstrated anti-tumor effects *in vivo* in the 22RV1 CRPC model. Our results prompt further translational and clinical studies with imipridones ONC201 or ONC201 in combination with enzalutamide or darolutamide for treatment of castrate resistant advanced or metastatic prostate cancer.

## Introduction

Advanced metastatic castrate resistant prostate cancer remains a major challenge as far as effective therapeutics especially among elderly patients with bone or organ metastases [1, 2]. While most advanced prostate cancer is androgen-sensitive and treatable with androgen deprivation, therapeutic resistance occurs [3, 4]. In the last 10-15 years, targeting androgen receptor signaling has been possible with AR pathway signaling inhibitors such as enzalutamide and darolutamide or androgen synthesis inhibition such as with abiraterone [5–7]. A subset of patients with prostate cancer have underlying molecular pathology involving loss of BRCA2 or other homologous recombination defects and those patients can be treated with PARP inhibitors as well as combinations that include PARP inhibition [8]. Other molecular defects such as PTEN loss or mutation, PIK3CA or Akt1/2/3 mutation have therapeutics under development. Gene fusions such as the ETS gene fusion TMPRSS2-ERG and mutations on p53, Rb, SPOP, CDK12 deletion and microsatellite instability have been described [9, 10]. Castrate-resistant prostate cancer becomes refractory to therapeutics, including a major subset that undergoes neuroendocrine differentiation, and these are in particular need of novel therapeutic strategies [11–16].

Our laboratory previously discovered and reported TRAIL-Inducing Compound #10 (TIC10) as a first-in-class anticancer agent [17]. The drug (imipridone TIC10/ONC201) advanced to first-in-human clinical trials in solid tumors as well as brain tumors through Oncoceutics, Inc. [18] and more recently through Chimerix, Inc. [19, 20] that is conducting trials including a large phase III trial (ACTION study) in H3K27M-mutated diffuse midline glioma [21], as well as early phase trials with ONC201 imipridone analogue ONC206 in brain tumors (NCT04732065, NCT04541082).

The first-in-human study of ONC201 conducted by Stein *et al*. observed that a number of patients with advanced prostate cancer with bone or soft tissue metastases experienced reduction in tumor size and improvement in symptoms [18, 22]. Some of the patients with prostate cancer were over the age of 90. Of 8 patients with advance prostate cancer who were treated with ONC201, 5 remained on therapy for greater than 6 months indicating an early signal in this disease. We previously discovered stimulation of natural killer cell activation by ONC201 [23]. A follow-up to the first-in-human ONC201 solid tumor study showed an association with immune markers and prolonged disease-free survival in patients with solid tumors who were treated with ONC201 [22]. To date, there has not been a phase II study of ONC201, its analogues, or combinations with other cancer therapies in the treatment of prostate cancer.

We previously explored combination treatments with ONC201 such as everolimus or enzalutamide [24]. Although the combination of ONC201 and everolimus showed potent synergy including *in vivo*, there was little interest to move this combination to the clinic as mTOR inhibitors such as everolimus have not shown clinical activity in prostate cancer. As ONC201 has shown an early signal in non-H3K27M-mutated non-brain tumors such as neuroendocrine tumors (pheochromocytomas, paragangliomas) [25], and since neuroendocrine differentiation is a feature of castrate-resistant advanced prostate cancer, this was another reason we were interested in exploring ONC201 and combinations with enzalutamide or darolutamide in prostate cancer. We had not previously conducted *in vivo* studies with ONC201 and enzalutamide and had not previously published studies of the combination of ONC201 or any imipridone plus darlolutamide in cell culture or *in vivo* [24].

In the present manuscript we further investigated combinations of ONC201 with enzalutamide as well as darolutamide including immunomodulatory and *in vivo* therapeutic effects in mice. We focused on darolutamide due to more favorable CNS toxicity profile for combination therapy. Our results support the further clinical translation of imipridones such as ONC201or ONC206 in combination with enzalutamide or darolutamide.

### Material and Methods Cell culture and reagents

Human prostate cancer cells lines, LNCaP, 22RV1, DU145 and PC3 were obtained from the American Type Culture Collection (ATCC). The cell lines are maintained in RPMI-1640 medium with L-glutamine (HyClone, Cytiva Life Sciences) with an addition of 10% FBS and 1% penicillin–streptomycin solution and were routinely essayed for mycoplasma contamination. ONC201 and ONC206 were obtained from Chimerix, Inc. Enzalutamide was purchased from MedKoo Biosciences (Cat. no. 201821). Darolutamide was purchased from MedKoo Biosciences (Cat. no. 206514). Apalutamide was purchased from Selleck Chemicals (Cat. no. S2840). DHT was purchased from ApexBio (Cat. no. S8214). The compounds were dissolved in DMSO at 20mmol/mL concentration and stored at -20°C.

### CellTiter-Glo^®^ Luminescent Cell Viability Assay

The CellTiter-Glo^®^ (CTG) reagent is obtained from Promega and stored in -80°C to preserve maximum light signal and other functional performances including half-life, linearity, and sensitivity. The CTG reagent was thawed at 4°C overnight or in a 22°C water bath and mixed gently before use per manufacturer’s instructions (Promega). A total of 5000 prostate cancer cells per well were seeded in an opaque-walled 96-well plate. After incubation for 24 hours at 37°C, cells were treated with ONC201 (0.313, 0.625, 1.25, 2.5, 5 and 10 μmol/L), ONC206 (0.313, 0.625, 1.25, 2.5, 5 μmol/L), enzalutamide (0.313, 0.625, 1.25, 2.5, 5, 10, 20 and 40 or 80 μmol/L), darolutamide (0.313, 0.625, 1.25, 2.5, 5, 10, and 20, 40 or 80 μmol/L), or apalutamide (0.313, 0.625, 1.25, 2.5, 5, 10, 20, 40 and 80 μmol/L), as single agents or in combinations. Matched volumes of medium without drug additions were added as vehicle control. After 72 hours of incubation at 37°C, the viability of cells was measured. 20 μl of CTG reagent was added to each well containing 100 μl of cell and medium mixture. The content in the plate was mixed for at least 2 minutes on an orbital shaker. Luminescence was measured using the IVIS Xenogen 200 imaging system and cell viability was recorded. All single treatment groups and combinations were performed in triplicates. Results were reported as mean values ± SD of the percentage of cell viability. Dose response curves were generated and the half-maximum growth inhibitory concentration (IC50) for each single treatment group was calculated using GraphPad Prism version 9.3.0.

### Drug Combination and Synergy Data Analysis

Combination indices were calculated using the CompuSyn software (ComboSyn Inc). Heat maps showing the combination indices (CI) were made by excel with combination indices indicating synergy highlighted in yellow. Heat maps showing synergy verses antagonism were generated using Combenefit software (Combenefit ^TM^). Data generated from combination assays were processed using the Highest Single Agent (HSA) classical synergy model as single experiments. A model-based combination dose-response surface based on the two single agent dose-response curves was generated to determine any additive or non-synergistic combination. Heat maps showing synergy and antagonism distribution was then developed by comparing the model-generated combination dose response surface to the experimental one. The sets of metrics provided by the Combenefit software were then used to interpret synergy or antagonism.

### Immunoblotting

Prostate cancer cells were regularly maintained in RPMI 1640 medium with L-glutamine (HyClone, Cytiva Life Sciences) supplemented with 10% FBS and 1% penicillin–streptomycin. A total of 2.5-3 x 10^5^ were seeded in medium and incubated overnight at 37°C with 5%CO2 before treatment. After treating, cells were washed with PBS and lysed with RIPA buffer (Cell Signaling) supplemented with one tablet of Complete^TM^ EDTA-free protease inhibitor cocktail (Sigma-Aldrich) and one tablet of Halt^TM^ phosphatase inhibitor (Thermo Fisher Scientific). The protein concentrations of the cell lysates were quantified using the BCA assay. To denature proteins, LDS sample buffer (Thermo Fisher Scientific) with 2% beta-mercaptoethanol was added and samples were placed on a heating block for 10 minutes. Equal amounts of proteins were then loaded into the 1.5 mm, 15 well NuPAGE 4-12% Bis-Tris gel (Thermo Fisher Scientific) and BLUEstain^TM^ Protein ladder (Goldbio) was used as a molecular weight marker. Gel was run on 1X MES running buffer [prepared from 20x MES (Thermo Fisher Scientific)] at 0.4 A for 40 minutes. Separated proteins were then transferred onto nitrocellulose membranes using wet transfer at 0.25 A for 1.5 hours. After blocking with 5% milk dissolved in TBS containing 0.1% Tween-20 for 30 minutes at room temperature, the membranes were incubated in primary antibodies diluted in 5% milk in TBS containing 0.1% Tween-20 (ATF4, cleaved-caspase 3 and PSA at 1:1000; cleaved-PARP at 1:2000; Ran at 1:5000) at 4°C overnight. After washing with 3 times with TTBS, membranes were incubated in appropriate secondary antibodies conjugated with horseradish peroxidase (goat anti-mouse IgG at 1:5000; goat anti- rabbit IgG at 1:10,000) for 1 hour at room temperature. Appropriate ECL reagents (Thermo Fisher Scientific) were added to the membranes for chemiluminescence detection using the Syngene Gel & Blot imaging system (SDI Group plc). The primary antibodies used in this study are as follows: antibodies against cleaved-PARP (Cell Signaling, cat. no. 9546s), AR (Cell Signaling, cat. no. 5153s), AR-V7 (Cell Signaling, cat. no. 68492s), cleaved caspase 3 (Cell Signaling, cat. no. 9661s), PSA/KLK3 (Cell Signaling, cat. no. 2474s), ATF4 (Cell Signaling, cat. no. 11815s) and Ran (BD Biosciences cat. no. BD 610341). The secondary antibodies used in this study were obtained from Thermo Fisher Scientific [goat anti-mouse IgG (H+L) 31430 and goat anti-rabbit IgG (H+L) 31460].

### Establishment of luciferase expressing human prostate cancer cell lines

Luciferase expressing prostate cancer cell line 22RV1-LUC was established to monitor mouse tumor growth by whole-body bioluminescence imaging using the IVIS Xenogen imaging system. 22RV1 prostate cancer cells were maintained at 60-70% confluency in RPMI 1640 medium with L-glutamine (HyClone, Cytiva Life Sciences) supplemented with 10% FBS and 1% penicillin–streptomycin. HEK293T cells were regularly maintained in DMEM/F12 1:1 medium with L-glutamine (Cytiva, cat. no. SH30023.01) supplemented with 10% FBS and ABX. A total of 6-7 x 10^6^ cells were seeded in medium and incubated overnight at 37°C with 5% CO2 and were maintained at 60-70% confluency prior to transfection. Transfection mix was made using psPAX2/Helper packaging plasmid (Addgene, cat. no. 12260), pMD2.G/VSVG.2 envelope plasmid (Addgene, cat. no. 12259) and shRNA/ pLX302 Luciferase-V5 puro (Addgene, cat. no. 47553), OptiMEM medium (Cellutron life Technologies, cat. no. 31985-070) and Lipofectamine 2000 (Life Technologies, cat. no. 11668- 027). HEK293T cells were incubated with the transfection mix first for 4-5 hours, then for 24 hours after addition of DMEM/RPMI medium with 15% and no Abx at 37°C with 5% CO2. Supernatants were collected and added to the 22RV1 cells directly with 8 µg/mL polybrene in equal amount of antibiotic-free medium. After incubation for 24 hours, cells were split and selected for bioluminescence expression.

### In vivo studies

All animal studies were approved and performed according to the Center for Animal Care Procedures (CARE) at Brown University and Institutional Animal Care and Use Committee (IACUC) protocols. Mouse experiments were performed free of pathogens. Four- to five-week- old male athymic sp/sp mice were purchased from Taconic and were quarantined in the institution’s animal facility for a week before any *in vivo* study was initiated. A total of 2x10^6^ luciferase expressing 22RV1 (22RV1-LUC) cells were suspended in 100 μl of ice-cold PBS and 100 μl of Matrigel® Basement Membrane Matrix (Corning^®^, cat. no. 354234). Anesthesia using 100 mg/kg ketamine was performed before a total of 200 μl cell suspension was inoculated subcutaneously into the rear left flank of the mice aged 6-8 weeks. The mice were weighed once a week to monitor signs of drug toxicity. Digital caliper measurements of the length (L), and width (W) of the tumors were taken twice a week to monitor tumor growth. When tumor volume reached an average of 150 mm^3^ to 200 mm^3^ (calculated using the formula: volume = ½ L x W^2^), mice were randomly assigned to control (vehicle) or indicated different treatment groups.

Vehicle was delivered in 40% PEG400, 5% Tween 80, 5% DMSO and 5% PBS. ONC201 (100 mg/kg, oral gavage, biweekly), enzalutamide (20mg/kg, oral gavage, daily) and darolutamide (50 mg/kg, oral gavage, twice daily) were delivered in a solution of 40% PEG400, 5% Tween 80, 5% DMSO and 5% PBS. Combination therapy experimental groups are the combination of ONC201 and enzalutamide or the combination of ONC201 and darolutamide. Whole-body bioluminescence imaging using the IVIS Xenogen imaging system was also carried out once a week. Mice were intraperitoneally injected with 15 mg/mL D-Luciferin (Goldbio, cat. no. LUCK-1G) and anesthetized with isoflurane. Signal intensities were recorded. The experiment was continued until the tumor burden had reached the ethically allowed limits at 2000 mm^3^ in volume or when the mouse had lost more than 20% of its initial body weight. At the end of the experiment, mice were sacrificed per Brown University CARE and IACUC protocols. Tumors and organs were harvested for flow cytometry, immunohistochemistry, serum cytokine profiling using a 42-plex murine cytokine panel for further analysis.

### Immunohistochemistry

Tumor and organ tissue sections were used for immunohistochemistry analysis. Resected 22RV1-LUC tumors and other organs from 5 mice each control and treatment groups were fixed in 10% formalin overnight, transferred into 70% ethanol and kept at 4°C before further paraffin treatment. Paraffin embedding, sectioning of slides, and H&E staining were performed by the Pathology Core Facility at Brown University. Sectioned tissue slides were deparaffined using xylene, rehydrated using decreasing concentrations of ethanol and TBS. Heat-induced antigen retrieval was conducted using a vegetable steamer (retrieval buffer: 10 mmol/L of sodium citrate buffer at pH = 6.0) for 20 minutes with temperature maintained at 95°C. After quenching with 3% hydrogen peroxide for 10 minutes and permeabilizing with TBST for 10 minutes, the samples were blocked using 2.5% horse serum (Vector Laboratories), before incubation with primary antibodies overnight at 4°C in a humid chamber. Horseradish peroxidase-conjugated secondary anti-rabbit antibody (Vector Laboratories, ImmPRESS^TM^, HRP-Horse Anti-Rabbit IgG Polymer secondary antibody, cat. no. MP-7401-15) diluted in blocking solution was added to the tissue slides and incubated in humid chamber for 40 minutes at room temperature. The slides were then developed with 3,3’-diaminobenzidine (Vector Laboratories, DAB/Ni Peroxidase Substrate Kit, cat. no. SK-4100) for 5 minutes and counterstained for 4 seconds using hematoxylin. After dehydrating with increasing concentrations of ethanol, the slides were mounted, and a cover slip was added. Images were taken.

### Flow cytometry

A total of 2-3 spleens and 2-3 tumors from each short term (1 week) treatment and control groups were processed for flow cytometry analysis. Mouse tumors were processed by finely cutting with razor blades before addition of digestion buffer (75U/mL Collagenase IV, 125 ug/mL Dispase II, 1% penicillin-streptomycin, in PBS). Tumor pieces were then placed in a 37°C water bath for 30 minutes until fully digested. Digested mouse tumors and spleens were then filtered through a 70 uM cell strainer and washed with PBS. Cells were isolated by centrifugation and resuspended in 2 mL RPMI-1640 medium (HyClone, Cytiva Life Sciences, with 10% FBS and 1% penicillin-streptomycin). 2 mL of 40% or 80% Ficoll-Paque PLUS (Cytiva, cat. no. 17-1440- 02, diluted in HBSS with 5% FBS) was added below the cell suspension. Immune cells were further isolated and collected on the interface after centrifugation at 2200 rpm for 30 minutes at room temperature. Processed tumor and spleen immune cells single cell suspensions were then resuspended in 1x Flow Cytometry Staining Buffer (R&D Systems). Staining for membrane surface proteins was conducted using conjugated primary antibody and antibody cocktail for 1 hour on ice. Antibody cocktail added to the samples include anti- CD45 SB600 (eBioscience, cat. no. 63-0451-82, Clone: 30-F11), anti-CD3 molecular complex (17A2) PE (BD Biosciences, cat. no. 555275), anti-CD335 (NKp46) (29A1.4) (Thermo Scientific, cat. no. 17-3351-82, Clone: PK136), anti-CD11b APC/Cy7 (BioLegend, cat. no. 101226, Clone M1/70), anti-KLRG1 PE/Cy7 (Thermo Scientific, cat. no. 25-5893-82, Clone: 2F1), and anti- CD253 (TRAIL) FITC (Miltenyi Biotec, cat. no. 130-102-447, Clone: N2B2). The controls for each tissue type include unstained control, viability only control stained with Zombie Violet^TM^ (BioLegend, cat. no. 423113), and single stained controls for each antibody used. Cells were then fixed using the EBioscience^TM^ IC Fixation buffer (Thermo Scientific) and incubated in the dark at room temperature for 30 minutes per manufacturer’s instructions. 200 μL of Flow Cytometry Staining Buffer (R&D Systems, cat. no. FC001) was added to the cells in 100 μL fixation buffer and filtered into flow cytometry tubes. All samples were kept at 4°C and analyzed via flow cytometry within 48 hours. All data was analyzed using a BD Biosciences LSR II Flow Cytometer and the FlowJo software version 10.8.1 (FlowJo, LLC). Gating strategies are as follows: NK cell: Live/NKp46+/KLRG1+/KLRG1-/TRAIL+/CD11b+.

### Cytokine profiling

Murine plasma samples were obtained and analyzed using an R&D Systems Murine Premixed 42plex Multi-Analyte Kit (R&D Systems, Inc.) and a Luminex 200 Instrument (LX200-XPON-RUO, Luminex Corporation) in accordance with the manufacturer’s instructions. Samples were analyzed for levels of GM-CSF, CXCL1/GRO alpha/KC/CINC-1, IL-7, GDF-15, CCL2/JE/MCP-1, IL-1 beta/IL-1F2, Chitinase-3-Like 1, VEGF, IL-2, IL-4, VEGFR2/KDR/Flk- 1, IL-6, IL-10, IL-17/IL-17A, IFN-gamma, IL-3, IL-16, CXCL10/IP-10/CRG-2, CCL5/RANTES, CCL7/MCP-3/MARC, CCL12/MCP-5, IL-33, Prolactin, M-CSF, CCL3/MIP-1alpha, IL-1 alpha/IL-1F1, CCL20/MIP-3alpha, TWEAK/TNFSF12, CXCL12/SDF-1 alpha, IGF-1, IL-27, BAFF/Blys/TNFSF13B, MMP-8, MMP-3, Granzyme B, Angiopoietin-2, CCL21/6Ckine, MMP-12, CCL12/Eotaxin, CCL22/MDC, and Dkk-1, values were reported in pg/mL.

### Statistical Analysis

For *in vitro* studies, all statistical analyses and graph generations were conducted using GraphPad Prism version 9.3.0. Data are presented as mean ± SD from at least three replicates for each experiment. The statistical significance comparing two groups was determined using the Student’s t-test. A minimal level of significance was set to be p < 0.05 (* p < 0.05, ** p < 0.01, *** p < 0.001). For *in vivo* studies, all statistical analysis and graph generations were conducted using GraphPad Prism version 9.3.0. Data are presented as mean ± SD from at least three technical replicates for each experiment. The statistical differences between treatment groups and controls were determined using one-way ANOVA test with a minimal level of significance of p < 0.05 (* p < 0.05, ** p < 0.01, *** p < 0.001).

## Results

### Darolutamide is efficacious in prostate cancer cell lines as single agent

It has been previously shown that darolutamide does not activate mutant AR such as AR(W742L), AR[T878A] and AR(F877L), and has a low inhibition constant (Ki) and maximal inhibitory concentration (IC50) when tested in AR-HEK293 cells stably expressing full-length AR [26]. When compared to other next-generation non-steroidal antiandrogens such as enzalutamide or apalutamide, darolutamide has a negligible blood-brain-barrier penetration, which may contribute to an improved CNS adverse effect (AE) [27]. To evaluate the *in vitro* efficacy of darolutamide against castration-resistant prostate cancer, 22RV1, LNCaP, PC3, and DU145 prostate cancer cell lines with distinct properties were treated with different concentrations of darolutamide in a CellTiter-Glo cell viability assay (**Table 1**).

**Table 1.** Prostate cancer cell lines used in the study.

| Cell Lines | Source of origin | AR status | Known key mutations | Castration sensitivity | Source | Reference |
| --- | --- | --- | --- | --- | --- | --- |
| **22RV1** | Primary xenograft | + | BRCA2 deficient, AR-V7 positive | Resistant | ATCC | [28] |
| **LNCaP** | Lymph node metastasis | + | DDR deficient | Sensitive | ATCC | [29] |
| **DU145** | Brain metastasis | - | BRCA (non-pathogenic) | Resistant | ATCC | [30] |
| **PC3** | Bone metastasis | - | PTEN deletion (homozygous) | Resistant | ATCC | [31] |

The half-maximal inhibitory concentration (IC50) was calculated for each cell line from the dose-response curves (**Figure 1**) to measure the potency of darolutamide in inhibiting cell proliferation in all four cell lines. All four cell lines displayed high sensitivity to darolutamide, with an IC50 of 46.6 µM for 22RV1, 33.8 µM for LNCaP, 32.3 µM for PC3 and 11.0 µM for DU145. Increased ATF4 expression was observed in darolutamide- treated 22RV1 and LNCaP cells and reduced prostate specific antigen (PSA) was observed in LNCaP cells (**Figure 2, 3**). Although co-treatment with DHT rescued some of the effects of darolutamide in reducing PSA and induction of ATF4, darolutamide showed better PSA reduction in LNCaP cells and ATF4 induction in both LNCaP and compared to cells treated with enzalutamide (**Figure 3**). Interestingly, darolutamide demonstrated higher inhibition potency in metastatic PC3 and DU145 cell lines, despite them being androgen independent. Western blot analysis was done to confirm the hypothesis that darolutamide may activate other pathways to reduce AR activity. In summary, darolutamide alone can upregulate the integrated stress response and reduce PSA level in castration-resistant or castration-sensitive prostate cancer *in vitro*.

**Figure 1.**
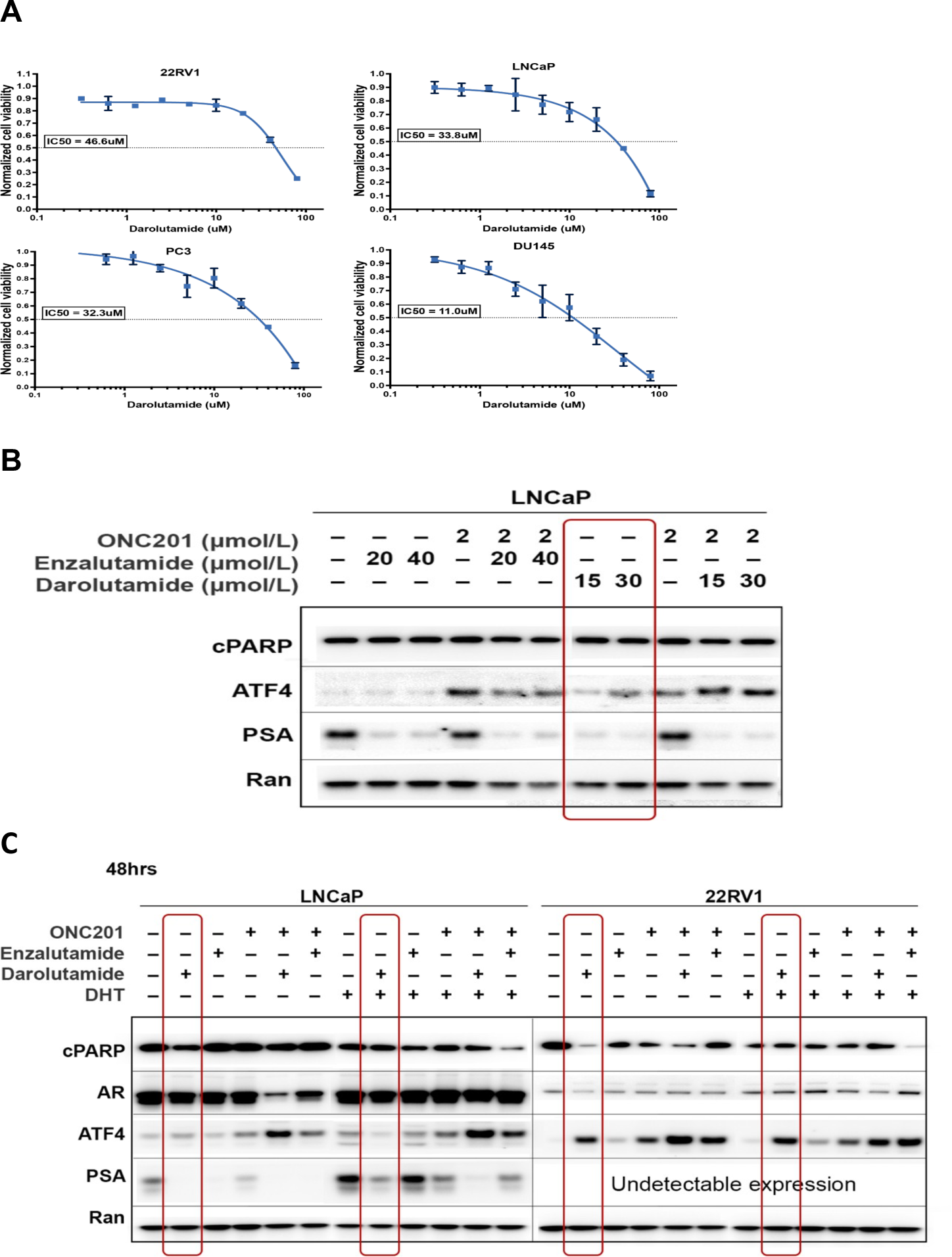
Dose-response curve and IC50s of darolutamide and effects on PSA, ATF4 in prostate cancer cell lines. **(A)** Proapoptotic and antiproliferation effects measured using the CTG assay in 22RV1, LNCaP, PC3 and DU145 cell lines 72 hours post treatment. Represented dose-response curves and calculated IC50s of darolutamide in each cell line are shown. **(B)** Darolutamide reduces PSA and induces ATF4 in LNCaP cell line. Western blot analysis of LNCaP cell line treated with ONC201, enzalutamide or darolutamide as single agents or in combinations. Red box highlights the single agent effect of darolutamide. **(C)** Darolutamide reduces PSA and induces ATF4 in LNCaP and 22RV1 cell lines regardless of DHT activation. AR signaling was stimulated by addition of 1µg/mL DHT. AR- expressing LNCaP and 22RV1 cell lines were co-treated with ONC201, enzalutamide or darolutamide as single agents or in combinations. Red box highlights the single agent effect of darolutamide in the presence and absence of DHT.

**Figure 2.**
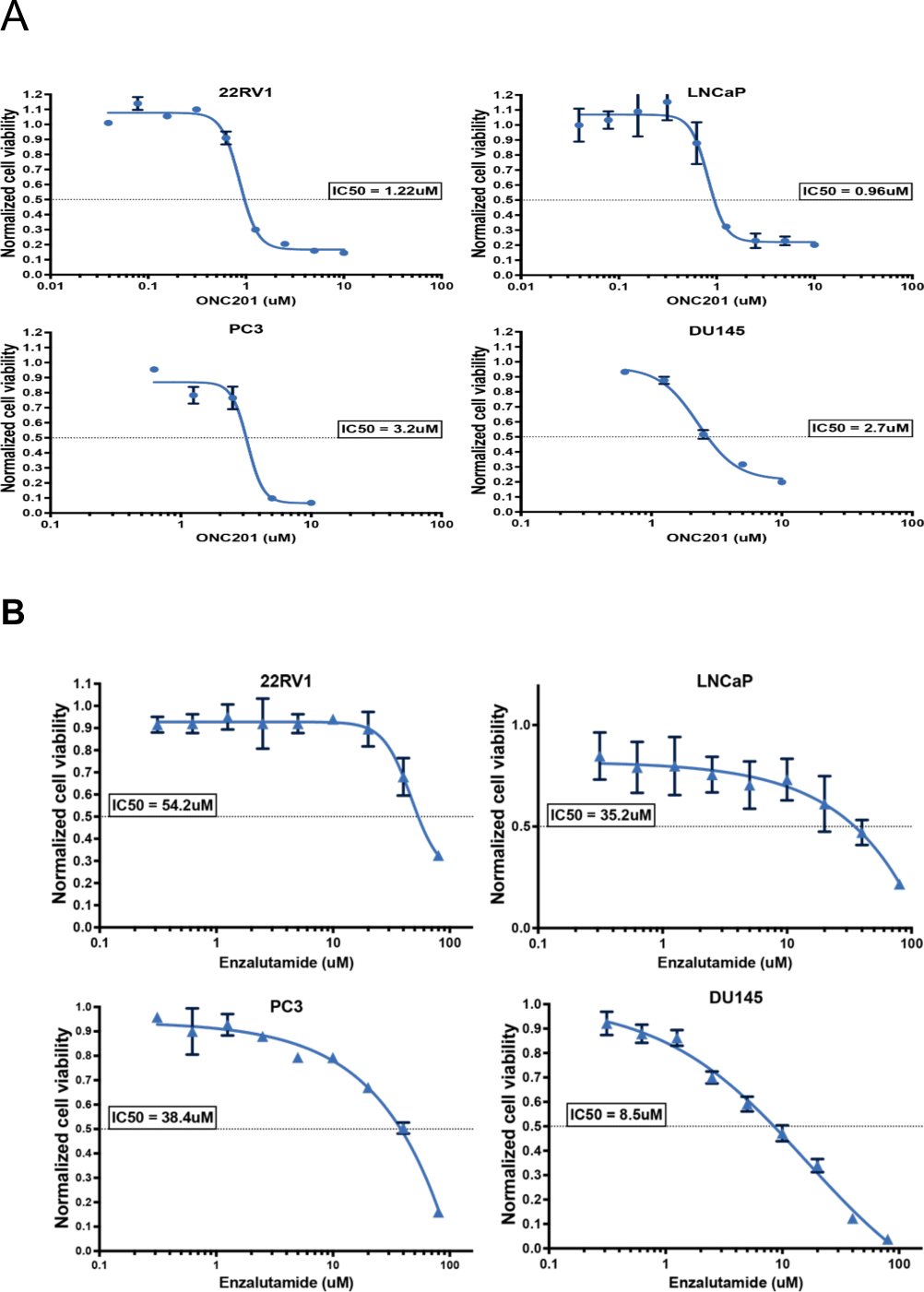
Representative dose-response curve and IC50s of ONC201 and enzalutamide in four prostate cancer cell lines. **(A).** Proapoptotic and antiproliferation effects measured using the CTG assay in 22RV1, LNCaP, PC3 and DU145 human prostate cancer cell lines 72 hours post treatment. Dose-response curves and calculated IC50s of ONC201 in each cell line are shown. (**B**) Proapoptotic and antiproliferation effects measured using the CTG assay in 22RV1, LNCaP, PC3 and DU145 human prostate cancer cell lines 72 hours post treatment. Represented dose-response curves and calculated IC50s of enzalutamide in each cell line are shown.

**Figure 3.**
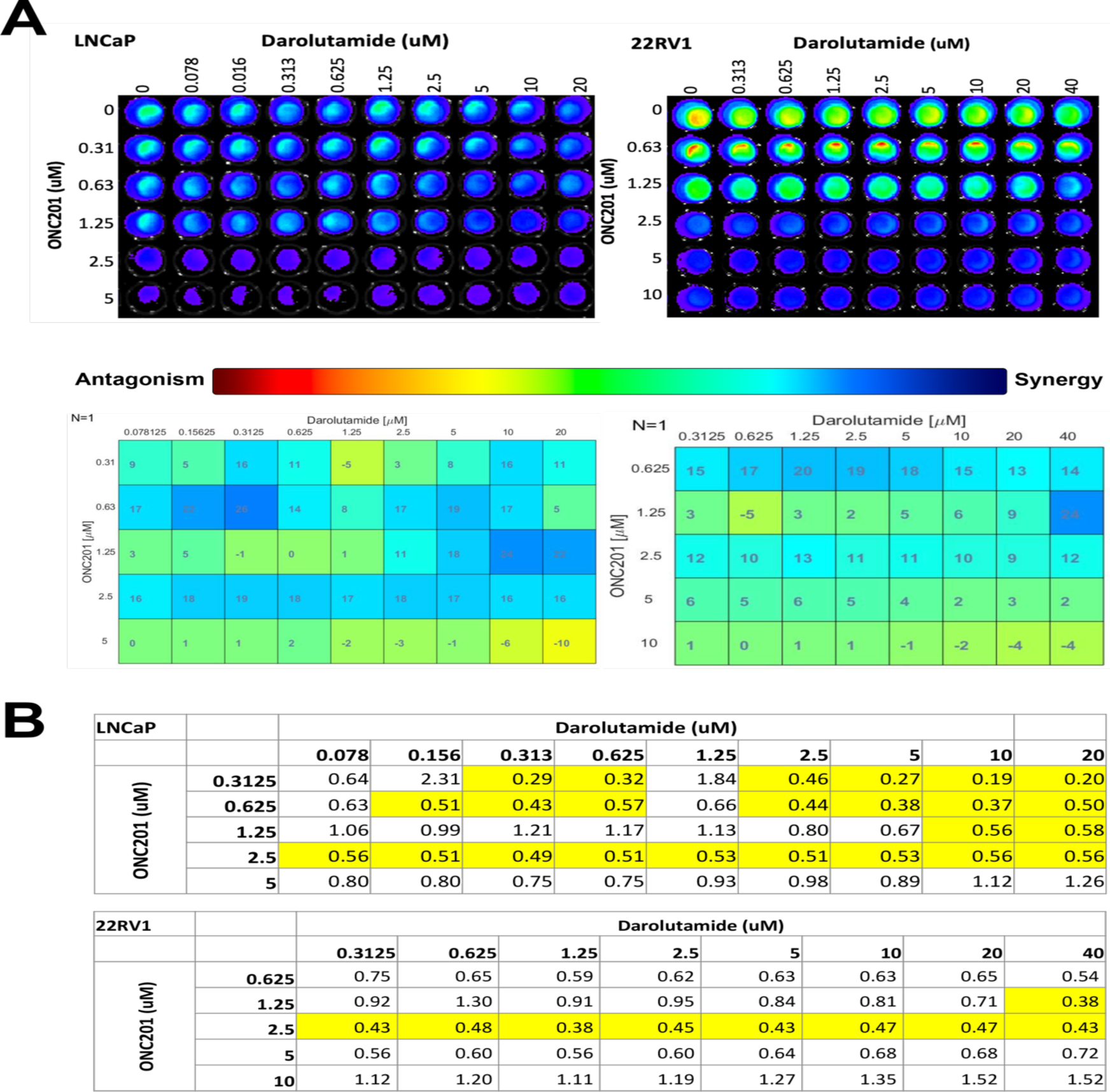
ONC201 synergizes with darolutamide against LNCaP and 22RV1 human prostate cancer cell lines *in vitro*. (A) Combinations of ONC201 (0-5umol/mL or 10umol/mL) and darolutamide (0-20umol/mL or 40mmol/mL) were evaluated in LNCaP and 22RV1 human prostate cancer cell lines 72hours post treatment using the CTG assay. Synergistic doses are shown**. (B)** Representative CI indices for combination of ONC201 and darolutamide in LNCaP and 22RV1 cell lines. Highlights indicate synergy.

### ONC201 synergizes with enzalutamide and darolutamide in prostate cancer *in vitro*

ONC201 has been previously reported to show anti-proliferative and pro-apoptotic efficacies in various prostate cancer cell lines by upregulation of TRAIL, DR5 and induction of the integrated stress response (ISR) through ATF4 [24]. With the observation that darolutamide is effective in treating prostate cancer as a single agent, its potential benefits in combination with ONC201 were evaluated. For better comparison, combination of ONC201 and enzalutamide was also included in the study. The IC50 of ONC201, enzalutamide and darolutamide were calculated from dose-response curves using CTG cell viability assay for single drugs and each indicated combination groups in LNCaP, 22RV1, PC3 and DU145 cells (**Figure 2**). A

Combination indices (CI) and heat maps showing synergy and antagonism in the ONC201 and enzalutamide or darolutamide groups in all four prostate cancer cell lines were generated to determine the synergistic effects (**Figure 3 – 6)**.

The androgen receptor (AR) signaling pathway is overactive in advanced forms of prostate cancer, altering expression of genes that contribute to uncontrolled cell proliferation and cancer survival [32]. Despite being hormone dependent, most forms of castration-resistant prostate cancer (CRPC) rely on AR and AR dependent signaling pathways and eventually lead to resistance to antiandrogen therapies [33]. Therefore, it was hypothesized that by combining ONC201 and androgen receptor antagonists, enzalutamide and darolutamide, the anti- proliferative and pro-apoptotic properties of ONC201 may be increased and subsequently enhance the inhibitory effect of enzalutamide and darolutamide against AR. Indeed, heat map results revealed that ONC201 synergized with darolutamide in LNCaP, 22RV1, PC3 and DU145 (**Figure 3A, 4A),** and enzalutamide in all of the four cell lines **(Figure 5A, 6A),** with the most synergistic combination doses in dark blue. Heat maps of CIs for each drug combinations also yielded consistent results showing best synergistic doses with combination indices lower than 0.5, highlighted in yellow **(Figure 3B, 4B, 5B, 6B)**.

**Figure 4.**
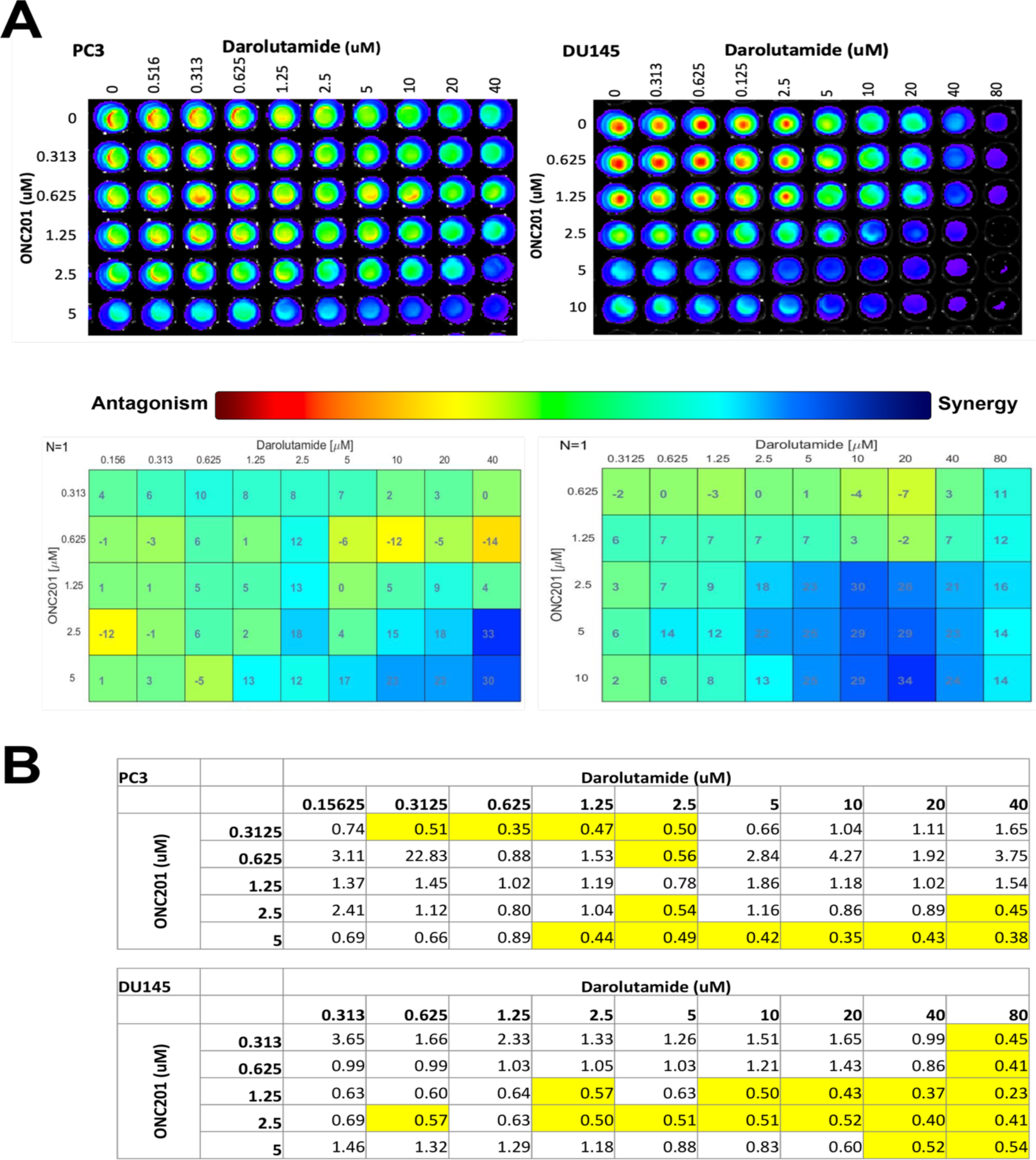
ONC201 synergizes with darolutamide against PC3 and DU145 human prostate cancer cell lines *in vitro.* **(A)** Combinations of ONC201 (0-5umol/mL or 10umol/mL) and darolutamide (0-40umol/mL or 80mmol/mL) were evaluated in PC3 and DU145 human prostate cancer cell lines 72hours post treatment using the CTG assay. Synergistic doses are shown. **(B)** Representative CI indices for combination of ONC201 and darolutamide in PC3 and DU145 cell lines. Highlights indicate synergy.

**Figure 5.**
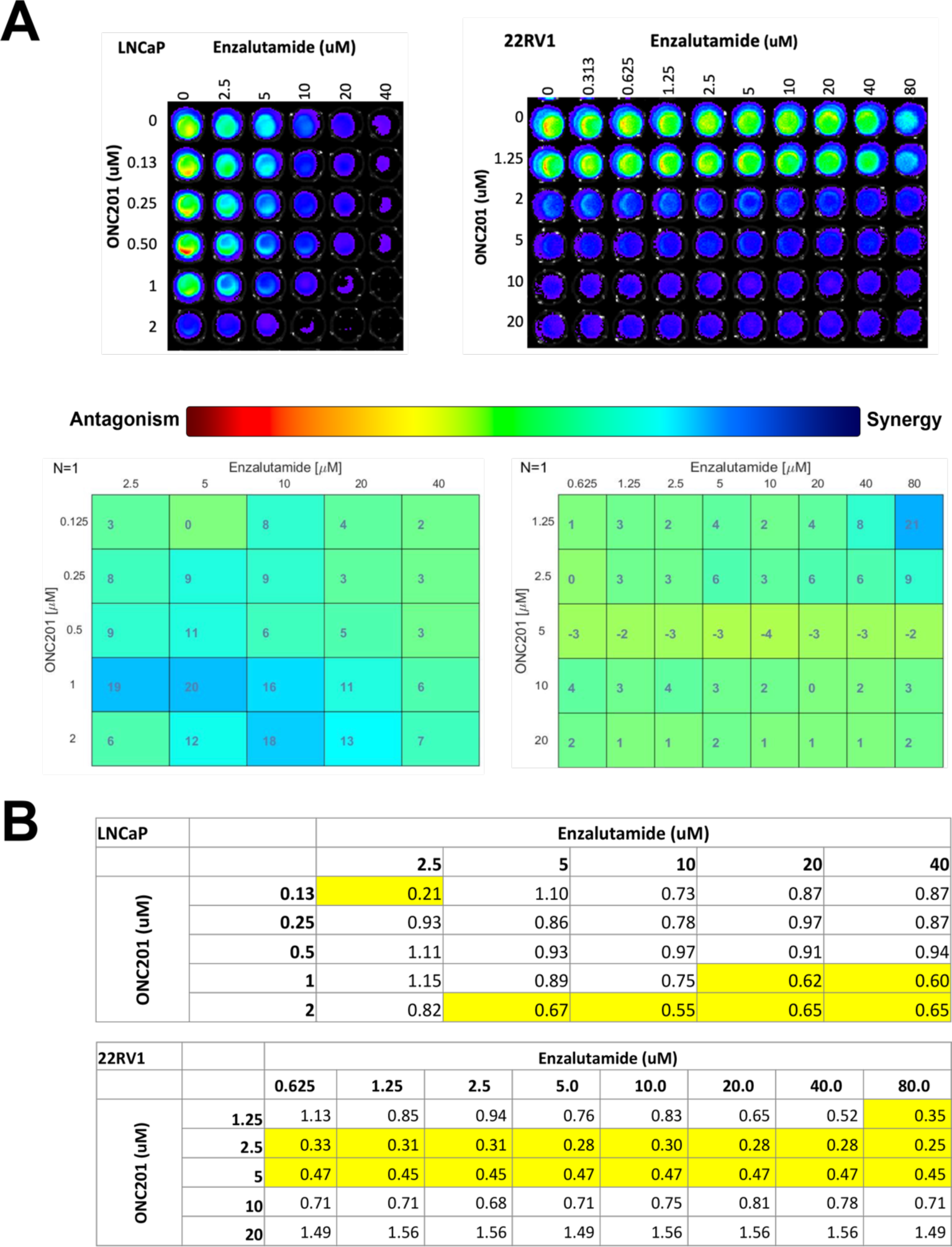
ONC201 synergizes with enzalutamide against LNCaP and 22RV1 human prostate cancer cell lines *in vitro.* **(A)** Combinations of ONC201 (0-2umol/mL or 20umol/mL) and enzalutamide (0-40umol/mL or 80mmol/mL) were evaluated in LNCaP or 22RV1 human prostate cancer cell lines 72hours post treatment using the CTG assay. Synergistic doses are shown. **(B)** Representative CI indices for combination of ONC201 and enzalutamide in LNCaP and 22RV1 cell lines. Highlights indicate synergy.

**Figure 6.**
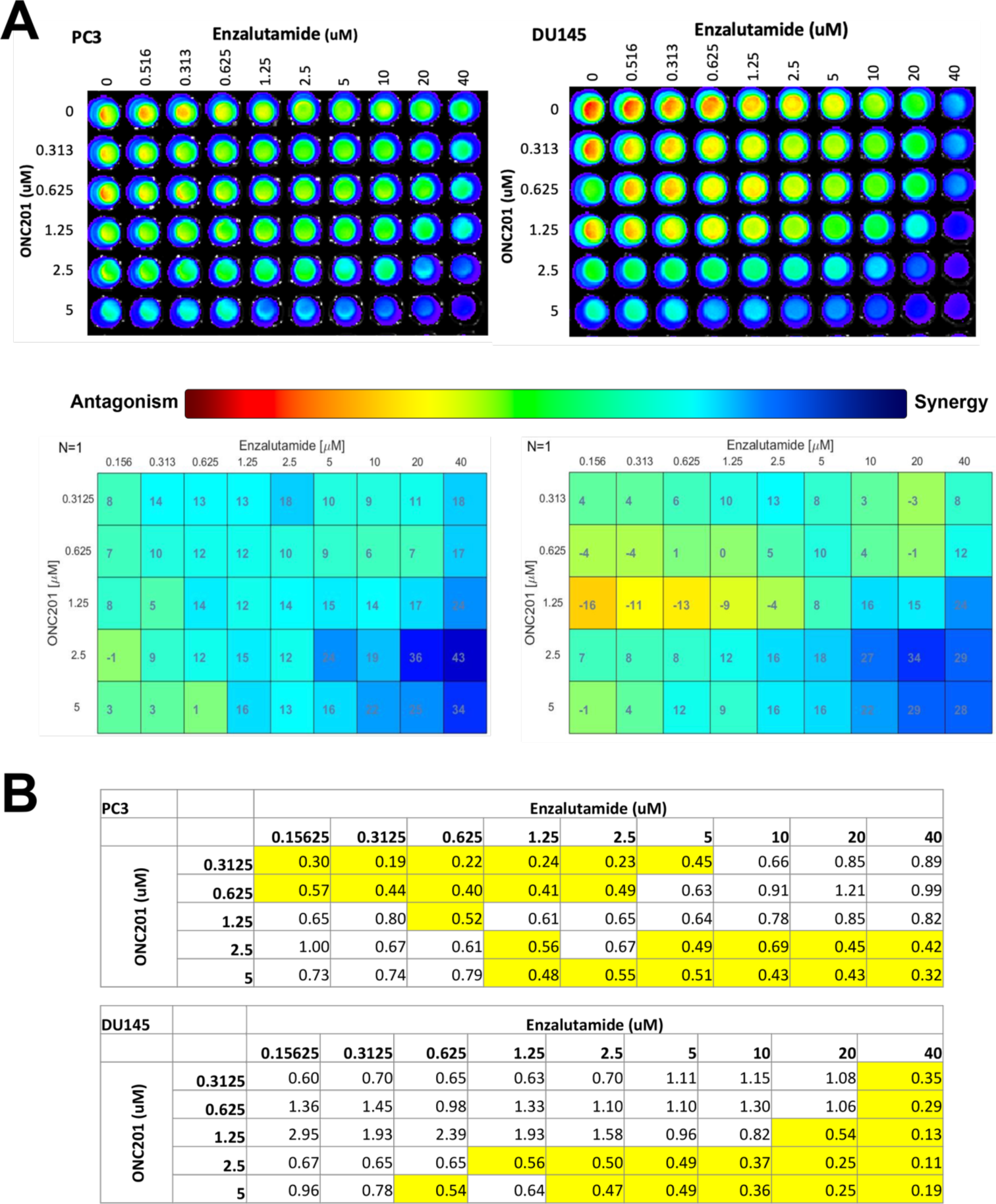
ONC201 synergizes with enzalutamide against PC3 and DU145 human prostate cancer cell lines *in vitro.* **(A)** Combinations of ONC201 (0-2umol/mL or 20umol/mL) and enzalutamide (0-40umol/mL or 80mmol/mL) were evaluated in LNCaP or 22RV1 human prostate cancer cell lines 72hours post treatment using the CTG assay. Synergistic doses are shown. **(B)** Representative CI indices for combination of ONC201 and enzalutamide in PC3 and DU145 cell lines. Highlights indicate synergy.

Cell viability assay results were evident that all four of the cell lines tested were sensitive to the combinations of ONC201 and enzalutamide or darolutamide at specific drug concentrations. Strong synergy of combination with ONC201 and darolutamide was observed across all four cell lines, whether metastatic (PC3, DU145, LNCaP) or non-metastatic (22RV1), castration resistant (22RV1, PC3, DU145) or sensitive (LNCaP), AR positive (22RV1, LNCaP) or negative (PC3, DU145) (**Figure 3, 4**). Darolutamide also required an overall lower synergistic dose when compared to the enzalutamide combination groups. This is consistent with the previously reported preclinical benefits of darolutamide, including having higher AR inhibitory potency and the ability to bypass activations of mutant AR, unlike the other antiandrogens such as enzalutamide or apalutamide [26]. The AR3 (AR-V7), a constitutively active AR variant whose transcriptional activities are independent of androgen or antiandrogen activities, has shown resistance against existing androgen receptor antagonists such as enzalutamide and apalutamide which may have led to the less synergistic combination of ONC201 and enzalutamide in AR-V7 positive 22RV1 cell line. Overall, the *in vitro* drug combination results showed improved antitumor efficacy with ONC201 and darolutamide or enzalutamide in treating both non-metastatic and metastatic prostate cancer.

### Combination of ONC201 and darolutamide reduce PSA level, activates ISR and promote apoptosis in prostate cancer cell lines

To further evaluate and compare the antitumor efficacies among the indicated treatment groups as well as between enzalutamide and darolutamide, LNCaP and 22RV1 cells were treated with ONC201, darolutamide or enzalutamide single agents or in combinations for 48 hours for protein marker analysis. The pro-apoptotic, anti-proliferation characteristics of ONC201 was confirmed in western blot analysis in 22RV1 cell line (**Figure 7A-B**), shown in PARP cleavage (cPARP) and cleaved caspase 3, which are markers for cells undergoing apoptosis.

**Figure 7.**
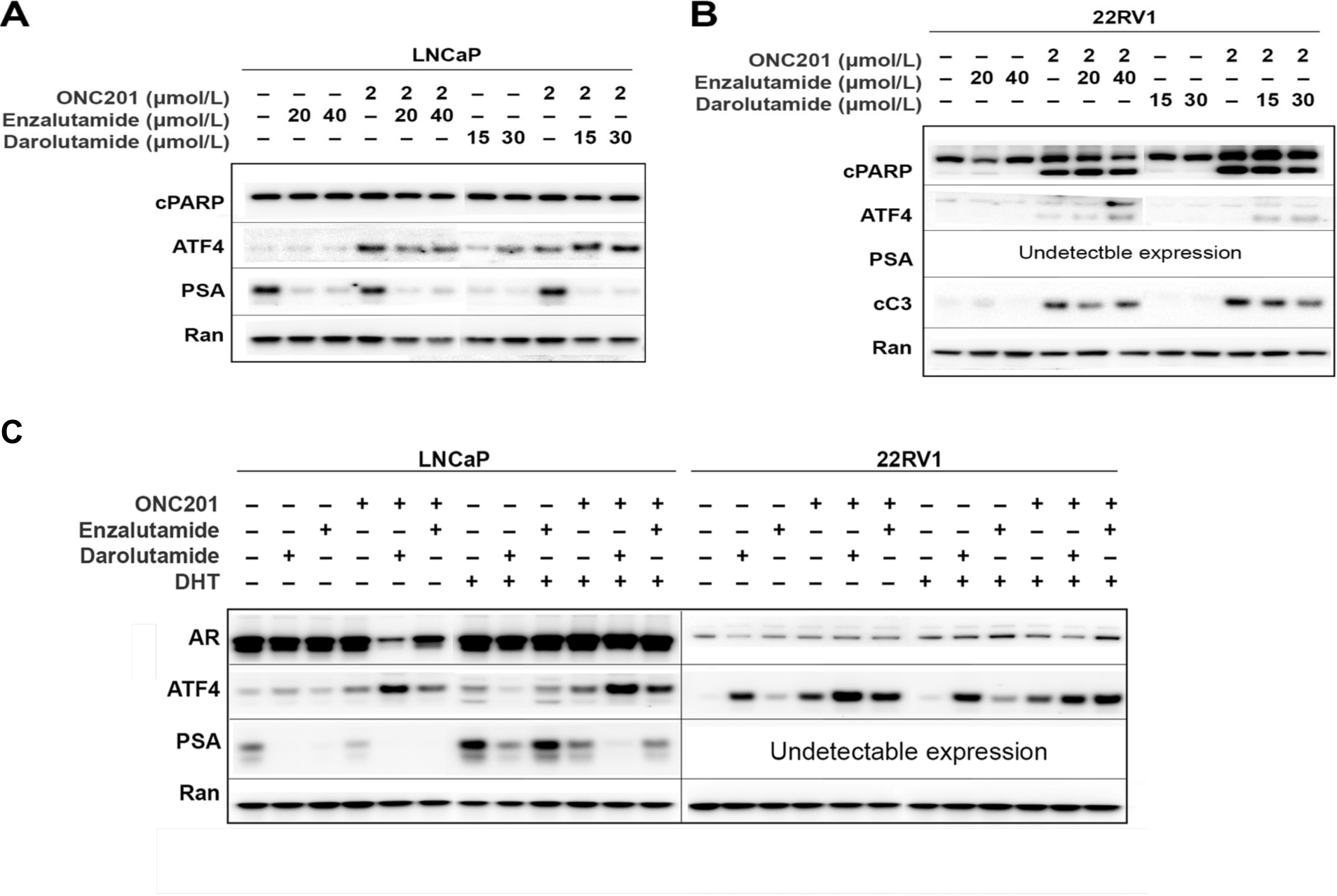
Effects of combinations of ONC201, darolutamide and enzalutamide on PSA, ATF4, PARP cleavage in human prostate cancer cell lines. **(A)** Western blot analysis of LNCaP cell line treated with 2umol/mL of ONC201, enzalutamide (20 umol/mL or 40umol/mL) or darolutamide (15 umol/mL or 30 umol/mL) as single agents or in combinations for 48 hours in comparison with untreated cells. **(B)** Western blot analysis of 22RV1 cell line treated with 2 umol/mL of ONC201, enzalutamide (20 umol/mL or 40 umol/mL) or darolutamide (15 umol/mL or 30 umol/mL) as single agents or in combinations for 48 hours in comparison with untreated cells. **(C)** AR signaling was induced by DHT (1 ug/mL) treatment in AR expressing LNCaP and 22RV1 cell lines. Western blot analysis of LNCaP and 22RV1 cells treated with 2umol/mL ONC201, 40 umol/mL enzalutamide or 30 umol/mL darolutamide as single agents or in combinations for 48 hours in comparison with untreated cells is shown.

It has been previously reported that ATF3, a stress response mediator downstream of ATF4 is a repressor of AR and is upregulated by ONC201, and the induction of both ATF4 and ATF3 is associated with reduction of PSA levels [24]. Western blot results at 48 hours showed upregulation of ATF4 when treated with ONC201, enzalutamide and darolutamide alone. Increased upregulation of ATF4 was observed in ONC201 and darolutamide combination groups, where higher dose of darolutamide yielded more ATF4 activation (**Figure 7A-B**). Though less ATF4 induction was seen in the 22RV1 cell line, similar results can be observed as the integrated stress response was induced in response to treatments with combinations of ONC201 and enzalutamide or darolutamide. PSA level was significantly reduced by enzalutamide or darolutamide alone or in combinations with ONC201 in LNCaP cell line. It is worth noting that ONC201 treatment alone showed less reduction in PSA level compared to darolutamide, enzalutamide and combinations, and ATF4 induction had no effect on PSA levels in LNCaP cell line.

Since darolutamide and enzalutamide demonstrated efficacies in reducing PSA levels, and combinations with ONC201 showed increased induction in ATF4, the effect of a more potent androgen receptor ligand, dihydrotestosterone (DHT), in activating AR and treatment efficacies was evaluated (**Figure 7C**). DHT (1 µg/mL) was co-treated with ONC201, enzalutamide, darolutamide or their indicated combinations. Western blot analysis revealed that darolutamide and enzalutamide alone reduced PSA level and induced ATF4 in both LNCaP and 22RV1 cell lines regardless of the presence or absence of DHT (**Figure 7C**). Combination of ONC201 and darolutamide also showed increased induction of AR in both tested cell lines when compared to combination of ONC201 and enzalutamide. In lane 5 of **Figure 7C**, combination of ONC201 and darolutamide significantly reduced AR expression, upregulated ATF4 and reduced PSA in LNCaP cell line. Although not as significant, in lane 11 of the same figure with an addition of DHT, the ONC201 and darolutamide combination had the same antiproliferation, pro-apoptotic and AR inhibition effects. Notably, DHT treatment did not increase AR expression in 22RV1 (**Figure 7C**). This is consistent with reports from previous studies suggesting that DHT is unable to stimulate the EGFR transcription as the AR isoforms in 22RV1 cells lack the COOH-terminal hormone binding domain (ARΔCTD) [34].

Taken together, the anti-proliferation and pro-apoptotic properties of ONC201 synergized with darolutamide, an antiandrogen with a higher AR inhibition potency than enzalutamide. The cell culture results shed light on the potential benefits of a novel combination therapy using ONC201 and darolutamide against CRPC. Results also support further evaluation of the antitumor efficacy, toxicity, and therapeutic benefits of the drug combinations *in vivo*.

### Trends of treatment-induced TRAIL activation within NK cells in the 22RV1 prostate cancer xenograft model

Since the combinations of ONC201 with enzalutamide or darolutamide were synergistic *in vitro,* both combinations were further investigated *in vivo* using a subcutaneously inoculated 22RV1-LUC xenograft model with immunodeficient male mice. Mice were randomly assigned to vehicle treated control groups or experimental groups and treated with 100 mg/kg ONC201 (oral gavage, twice a week), 20 mg/kg enzalutamide (oral gavage, daily), or 50 mg/kg darolutamide (oral gavage, twice daily) as single agents or combinations of ONC201 and enzalutamide or darolutamide. Tumor volumes were measured using a digital caliper twice a week. Mice were further divided into short term (1 week, n = 2 or 3 per group) and long term (6 weeks, n = 6 or 7 per group) treatment groups.

Immune modulations were evaluated in the short-term (1 week) treatment groups via flow cytometry analysis. Levels of spawning/homing (**Figure 8A**) and intratumor/migrating (**Figure 8B**) KLRG1-, TRAIL+, and CD11b+ NK cells were measured to determine stages of the NK cells, TRAIL activation, and NK cell maturation, respectively. Although no significance was shown due to sample size and the short-term treatment timeline, trends of intratumor NK cells were observed (**Figure 8B**), represented by the NKp46 + cells. Overall trends of increasing TRAIL induction by the antiandrogen and combination groups with ONC201 were also presented in the tumors (**Figure 8B**). Integrin CD11b is used as NK cell maturation marker in mice, with CD11b+ NK cells indicating maturation. In **Figure 8B**, trends of migrating NK cells can be seen in the tumors of the mice. NK cell maturation indicated by increasing levels of CD11b+ cells suggest treatment-induced immune response, which may contribute to eventual partial tumor regression.

**Figure 8.**
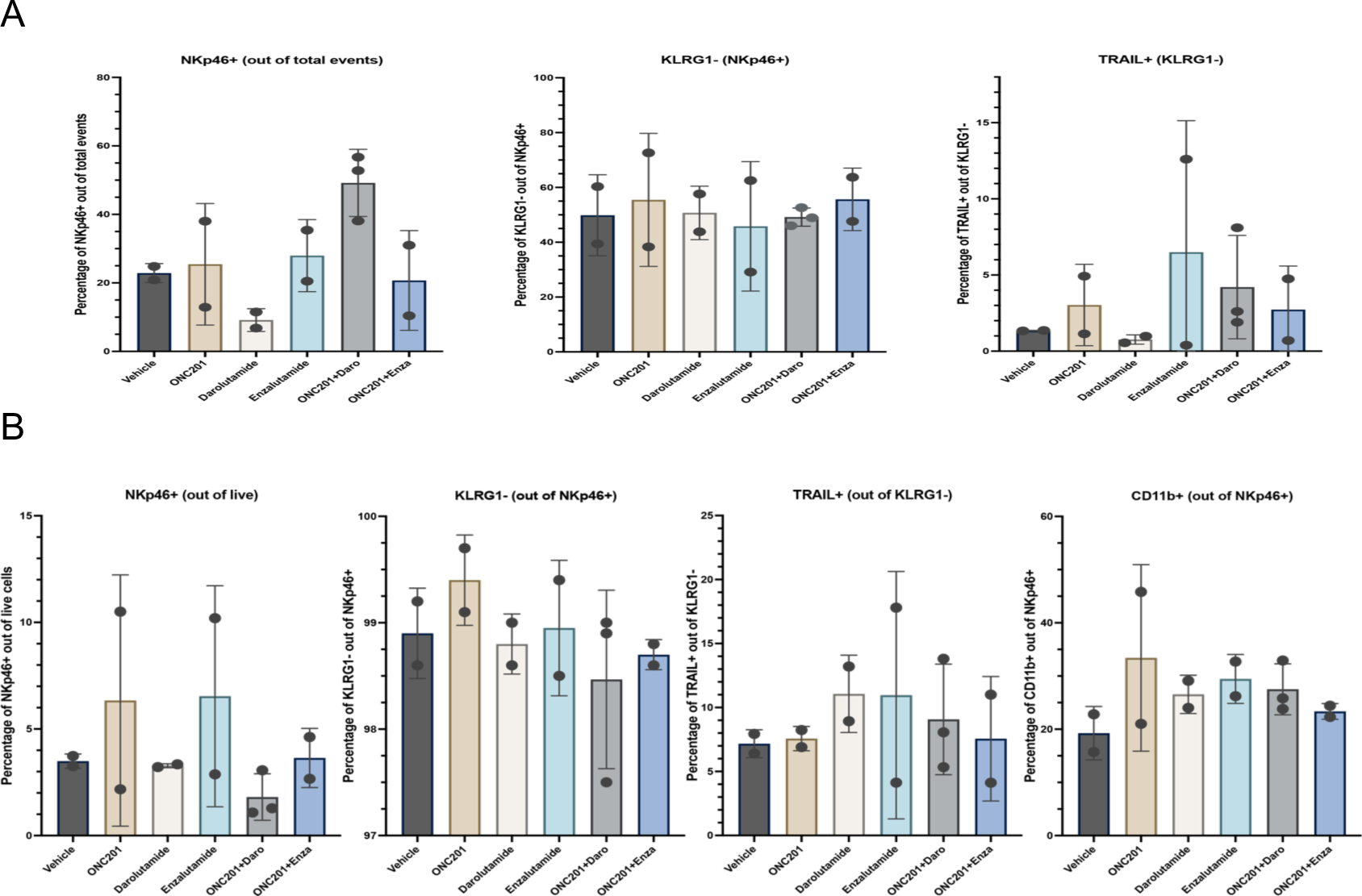
Mouse spleen and tumor flow cytometry results showing trends of TRAIL activation within NK cells. **(A)** Graphs summarizing flow cytometry data for KLRG1- and TRAIL+ NK cells in spleens of mice in indicated control and treatment groups 7 days post treatment. Results represented as mean ± SEM of one experiment, showing trends indicating circulating NK cells and TRAIL activation in treatment groups. (n = 2 for all experimental and control groups except for group treated with combinations of ONC201 and darolutamide where n = 3). **(B)** Graphs summarizing flow cytometry data for KLRG- TRAIL+ CD11b+ NK cells in 22RV1- LUC subcutaneous tumors of mice in indicated control and treatment groups 7 days post treatment. Results represented as mean ± SD of one experiment with 2 to 3 mice per group, showing trends indicating infiltrating NK cells and TRAIL activation in treatment groups.

### Treatment induced immune modulations may contribute to partial tumor regression in the 22RV1 prostate cancer xenograft model

Whole blood was collected from mice (n = 9 for partial responders, n = 21 for non- responders) upon sacrifice and purified serum was analyzed for a panel of murine cytokines (**Figure 9A**). The overall results were largely heterogenous across all cytokines when comparing single treatments, combinations, and control groups (**Figure 9A**). However, after regrouping as partial responders (mice that showed signs of tumor regression) and non- responders (mice that did not show signs of tumor regression), a comparison of treatment effect (mean values ± SD ) between the two groups yielded statistically significant increase in the following cytokines: GM-CSF (p = 0.0260), Chitinase-3-Like-1 (p = 0.0125), IL-2 (p = 0.0463), IFN-gamma (p = 0.0011), IL3 (p = 0.0098), CCL7/MCP-3/MARC (p = 0.0403), Prolactin (p = 0.0078), IL-1 alpha/IL-1F1 (0.0285), and MMP-12 (0.0472) (**Figure 9B** with red box highlights, **Figure 9C**). In **Figure 9D**, volcano plot also displayed trends towards statistical significance in upregulations of pro-inflammatory and pro-apoptotic cytokines include CXCL/1/GRO alpha/KC/CINC-1 in ONC201 treatment group (**Figure 9D subpanel A**), GDF-15, CCL7/MCP-3/MARC, CCL3/MIP-1alpha, CXCL1/GRO alpha/KC/CINC-1, IL-6 in enzalutamide group (**Figure 9D subpanel B**), IL-17/IL-17A, MMP-8, GDF-15, and TWEAK/TNFSF12 in darolutamide group (**Figure 9D subpanel C**), Dkk-1 in ONC201 and darolutamide combination group (**Figure 9D subpanel E**), and IL-16 and IL-6 in ONC201 and enzalutamide combination group (**Figure 9D subpanel D**). Downregulations of IGF-1 in all treatment groups was observed. Other cytokines that exhibited trends of downregulation across the treatment groups include BAFF/Blys/TNFSF13B and VEGFR2/KDR/Flk-1. All comparisons in the volcano plot were relative to control group.

**Figure 9.**
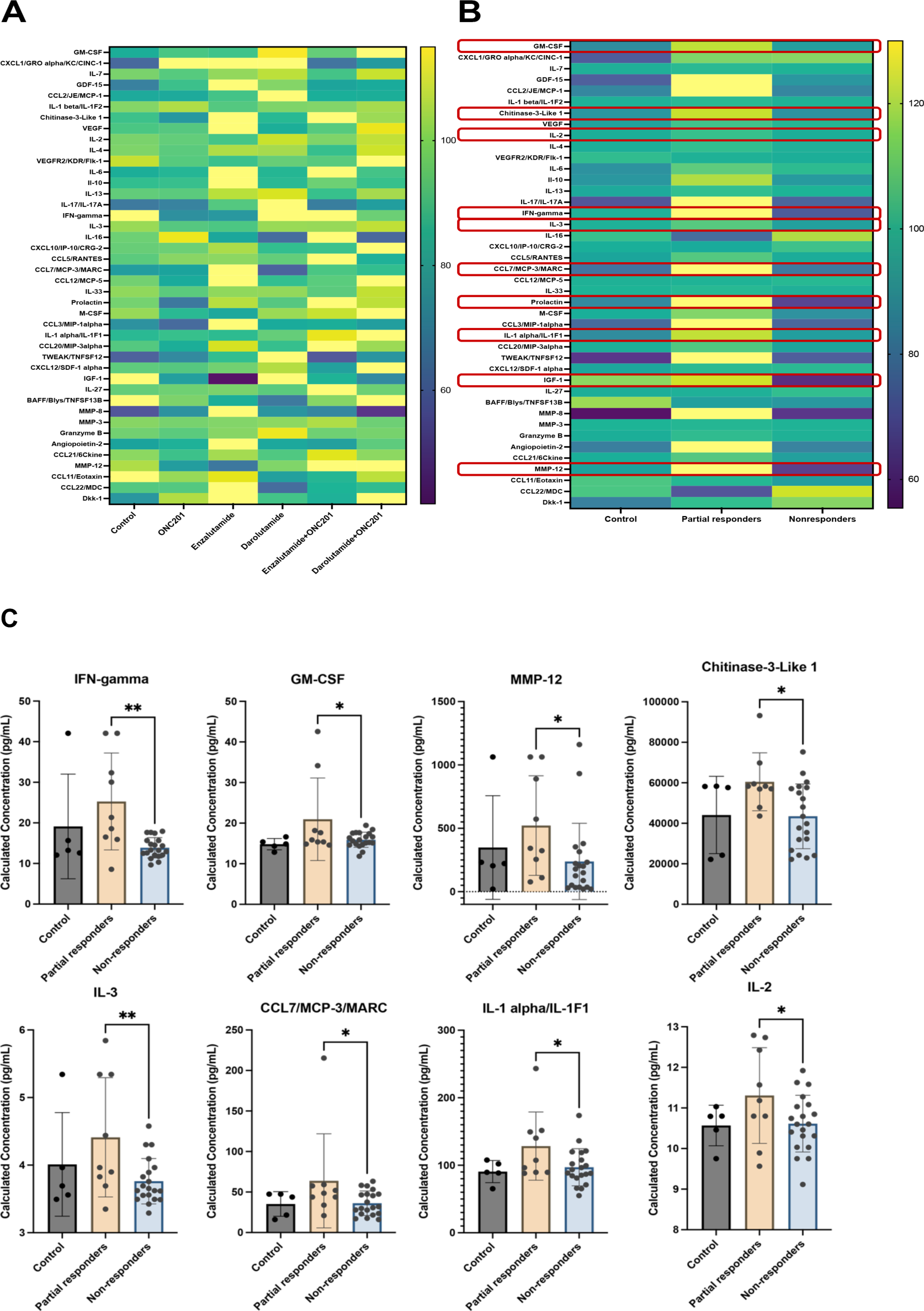
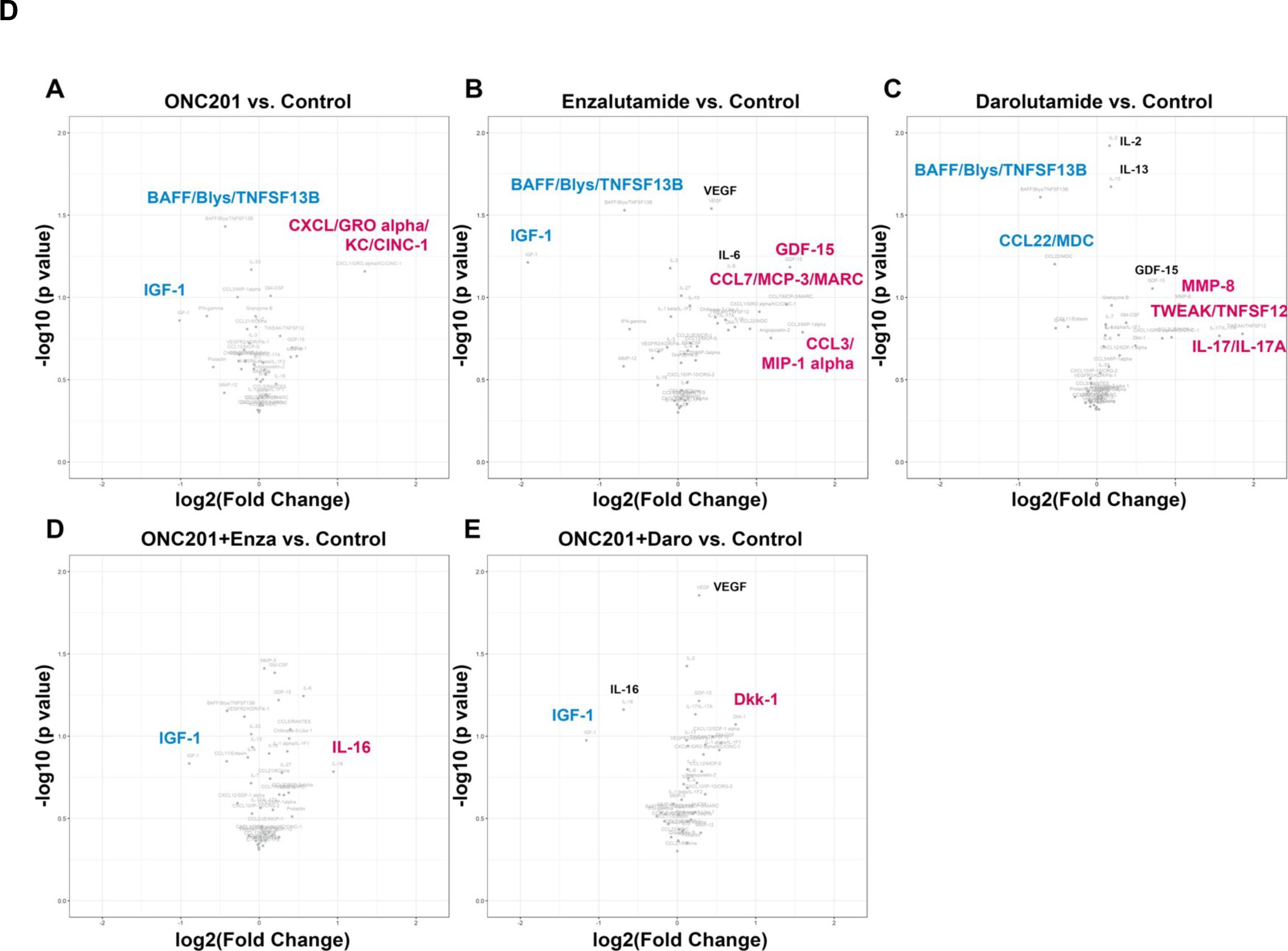
Cytokine profiling of serum in mice. **(A)** Heat map of cytokine expression levels for mice in control (vehicle) group or experimental groups treated with ONC201, enzalutamide, darolutamide or indicated combinations. The row maximum is shown in full yellow and row minimum is shown in full dark purple. **(B)** Heat map of relative cytokine expression levels for mice that showed partial tumor regression (partial responders) and those that did not show any significant regression (non-responders). The row maximum is shown in full yellow and row minimum is shown in full dark purple. Rows where statistically significant differences were calculated by one-way ANOVA test followed by pairwise comparisons are highlighted in red. (Results were normalized to the means within each cytokine sub-column and are shown in percentage. 0% was defined as = 0 and 100% was defined as the largest mean in each data set). (C) Blood serum was collected from mice when harvesting at the end of the experiments and analyzed using a 42- plex murine cytokine panel. Graph results of serum cytokine profiling of mice post treatment that show differential expression with statistical significance for partial responders to treatments verses non- responders. Analysis was done using the average of two technical replicates. Sample values that were above the upper limit of detection were recoded as the upper limit of detection; sample values that were below the lower limit of detection were recorded as 0 or a half of the lower limit of detection. * p < 0.05; ** p < 0.01, by one-way ANOVA. (D) Volcano plots illustrating the immune modulation of mouse serum cytokines. **(a)** Comparisons made between control and ONC201 treated groups. **(b)** Comparisons between enzalutamide and control groups. **(c)** Comparisons between darolutamide and control groups. **(d)** Comparisons between control group and the group treated with combination of ONC201 and enzalutamide. **(e)** Comparisons between control group and combination of ONC201 and darolutamide. Log2 (fold change) > 1 is indicated as upregulation (red), log2 (fold change) < 1 is indicated as downregulation (blue). Significance threshold used: p < 0.05.

Taken together, the upregulation of pro-inflammatory cytokines promoting immune responses and downregulation of cytokines that promote growth factor or cell survival pathways may have contributed to the partial tumor regression seen in the experimental groups.

### ONC206 shows strong synergy with enzalutamide, darolutamide or apalutamide in DU145 and PC3 prostate cancer cell lines

With previous experiments validating the antitumor efficacies of ONC201 and the combinations of ONC201 and second generation NSAAs, it was hypothesized that ONC206, a more potent analogue of ONC201 that has shown preclinical pro-apoptotic effects in endometrial cancer [18], would yield better synergy with the new generation agents against prostate cancer. Two metastatic castration-resistant prostate cancer cell lines, DU145 and PC3 (**Table 1**) were used and treated with ONC206 and three second generation antiandrogens, enzalutamide, darolutamide, apalutamide, as single agents or in combinations. Single agent dose-response curves were generated by the CTG viability assay, and IC50s for ONC206 (**Figure 10A**), enzalutamide (**Figure 2B**), darolutamide (**Figure 1**), and apalutamide (**Figure 10B**) were calculated. The CIs and heat maps indicating synergy or antagonism in the ONC206 and enzalutamide, darolutamide or apalutamide groups were generated to determine the synergistic effects (**Figure 10C**, **D**, **Table 2**).

**Figure 10.**
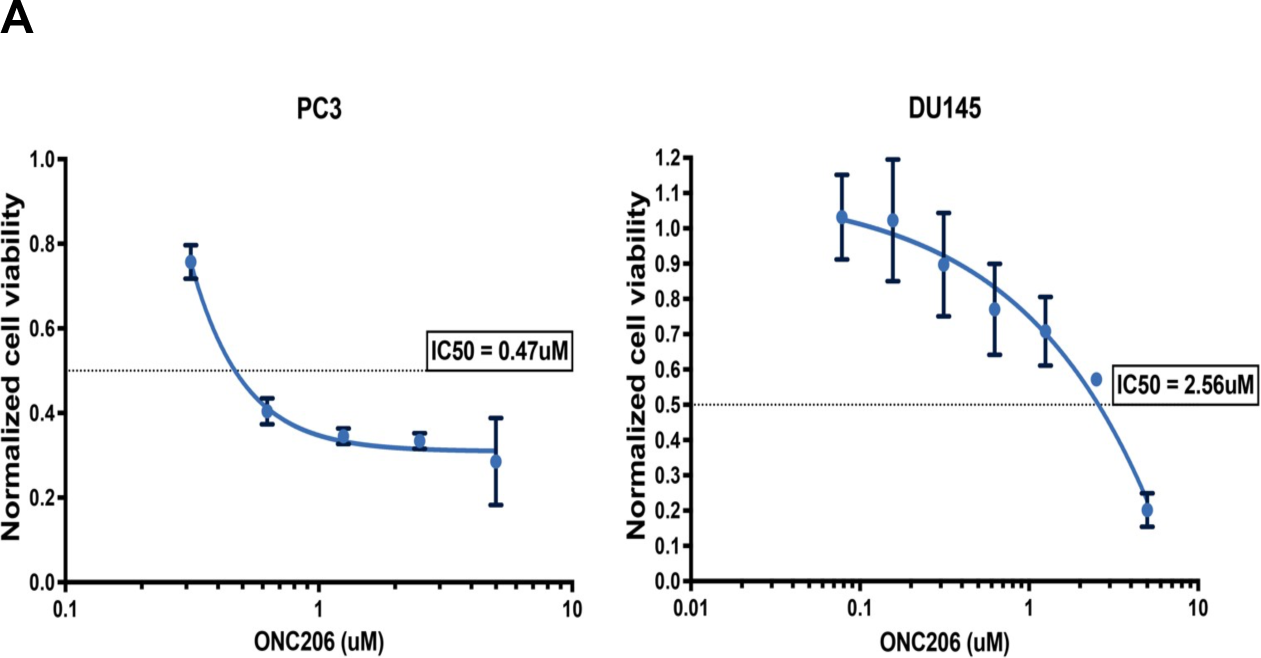
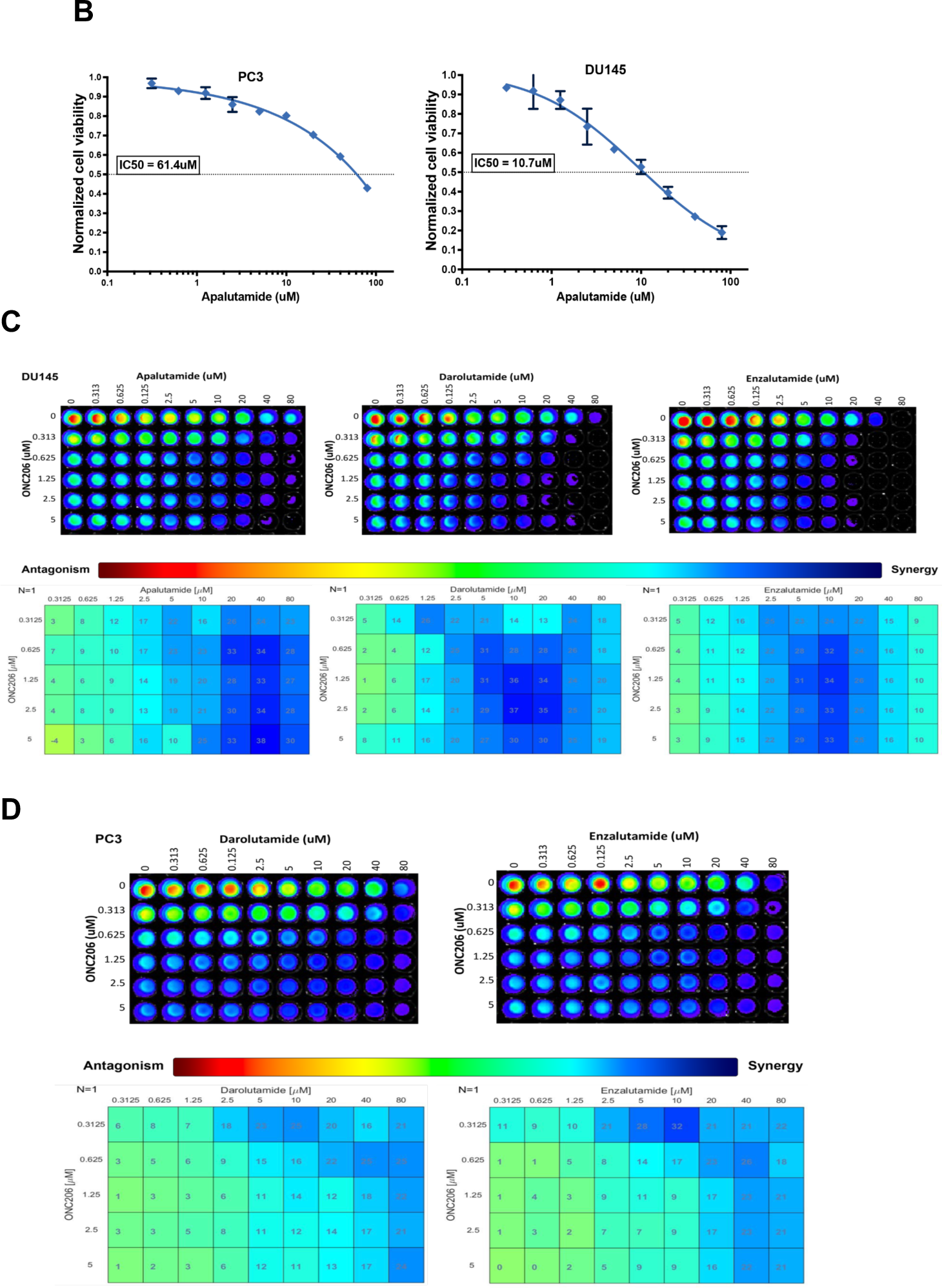
Representative dose-response curve and IC50s of ONC206 and apalutamide in two prostate cancer cell lines along with synergy analysis of ONC201 with enzalutamide, darolutamide and apalutamide. **(A)** Proapoptotic and antiproliferation effects measured using the CTG assay in PC3 and DU145 human prostate cancer cell lines 72 hours post treatment. Represented dose- response curves and calculated IC50s of ONC206 in each cell line are shown. **(B)** Proapoptotic and antiproliferation effects measured using the CTG assay in PC3 and DU145 human prostate cancer cell lines 72 hours post treatment. Represented dose- response curves and calculated IC50s of apalutamide in each cell line are shown. **(C)** Combinations of ONC206 (0-5 umol/mL), apalutamide (0-80 μmol/mL), darolutamide (0-80 μmol/mL) and enzalutamide (0-80 μmol/mL) were evaluated in DU145 human prostate cancer cell lines 72 hours post treatment using the CTG assay. Synergistic doses are shown. **(D)** Combinations of ONC206 (0-5 μmol/mL), darolutamide (0-80 μmol/mL) and enzalutamide (0-80 μmol/mL) were evaluated in PC3 human prostate cancer cell lines 72 hours post treatment using the CTG assay. Synergistic doses are shown.

**Table 2.**
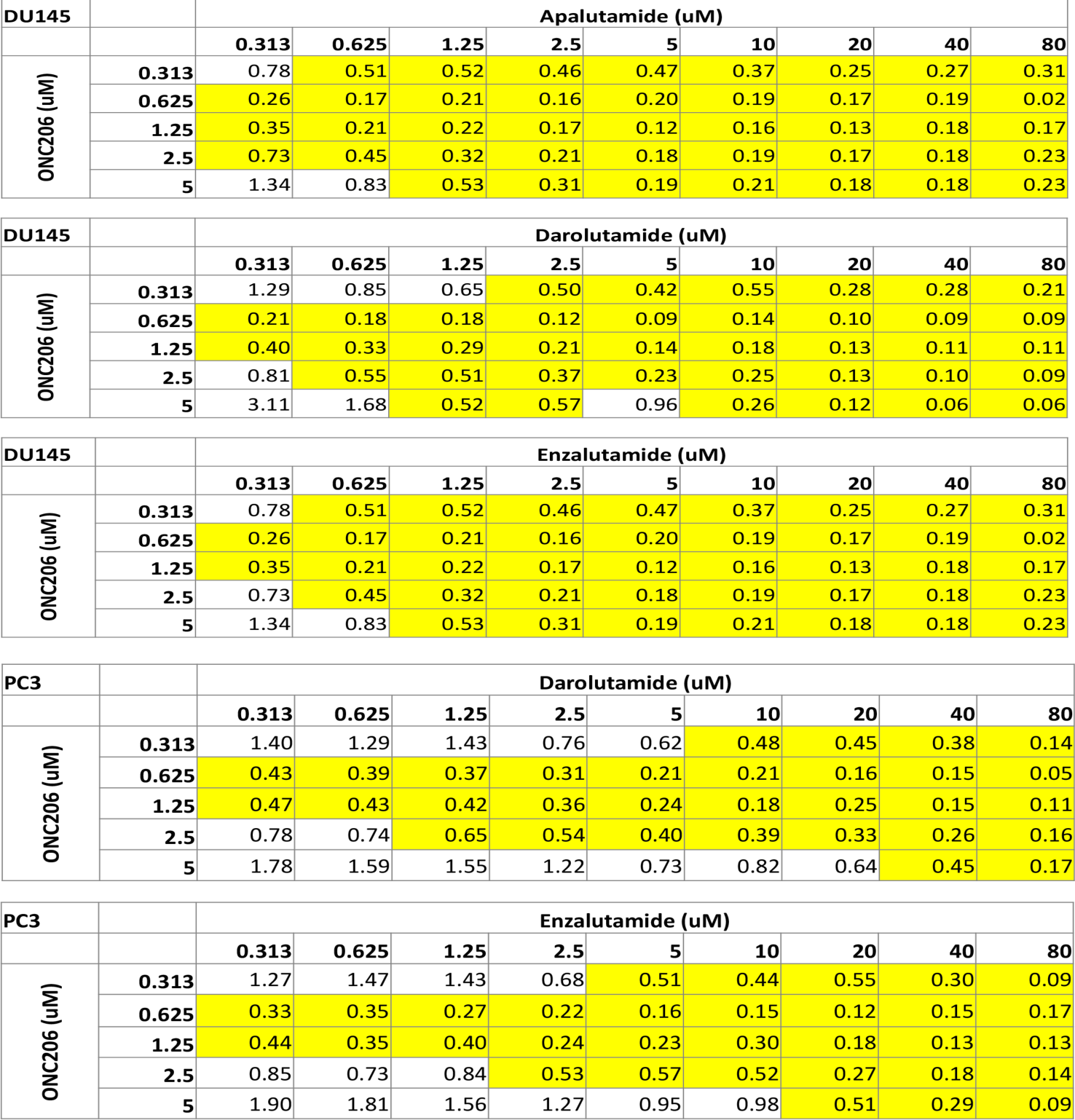
Representative CI indices for combinations of ONC206 and anti-androgen agents in DU145 and PC3 prostate cancer cell lines. Highlights indicate synergy.

*In vitro* results indicated that both PC3 and DU145 cell lines are sensitive to ONC206 and strong synergy was observed in both PC3 and DU145 cell lines treated with indicated combination groups (**Figure 10C, 10D**). An overall lower CI, shown in PC3 and DU145 (**Table 2**) proved the hypothesis that ONC206 yielded better synergistic effects when combined with antiandrogens in cell viability assay. The preliminary results with ONC206 support further studies using the indicated combination therapy *in vitro* and *in vivo* against advanced prostate cancer.

### Sensitivity of 22RV1 *in vivo* to ONC201 and ONC201 plus darolutamide

Treatment with ONC201 at 50 or 100 mg/kg 3x per week demonstrated anti-tumor efficacy at the higher dose (**Figure 11A**). Because 100 mg/kg ONC201 was highly effective as monotherapy, we reduced the dose to 75 mg/kg and observed anti-tumor efficacy with the combination of ONC201 plus darolutamide *in vivo* (**Figure 11B**) and this was associated with increased TRAIL expression (**Figure 11C**). We did not observe alterations in ATF4, AR, or Ki67 in residual tumors.

**Figure 11.**
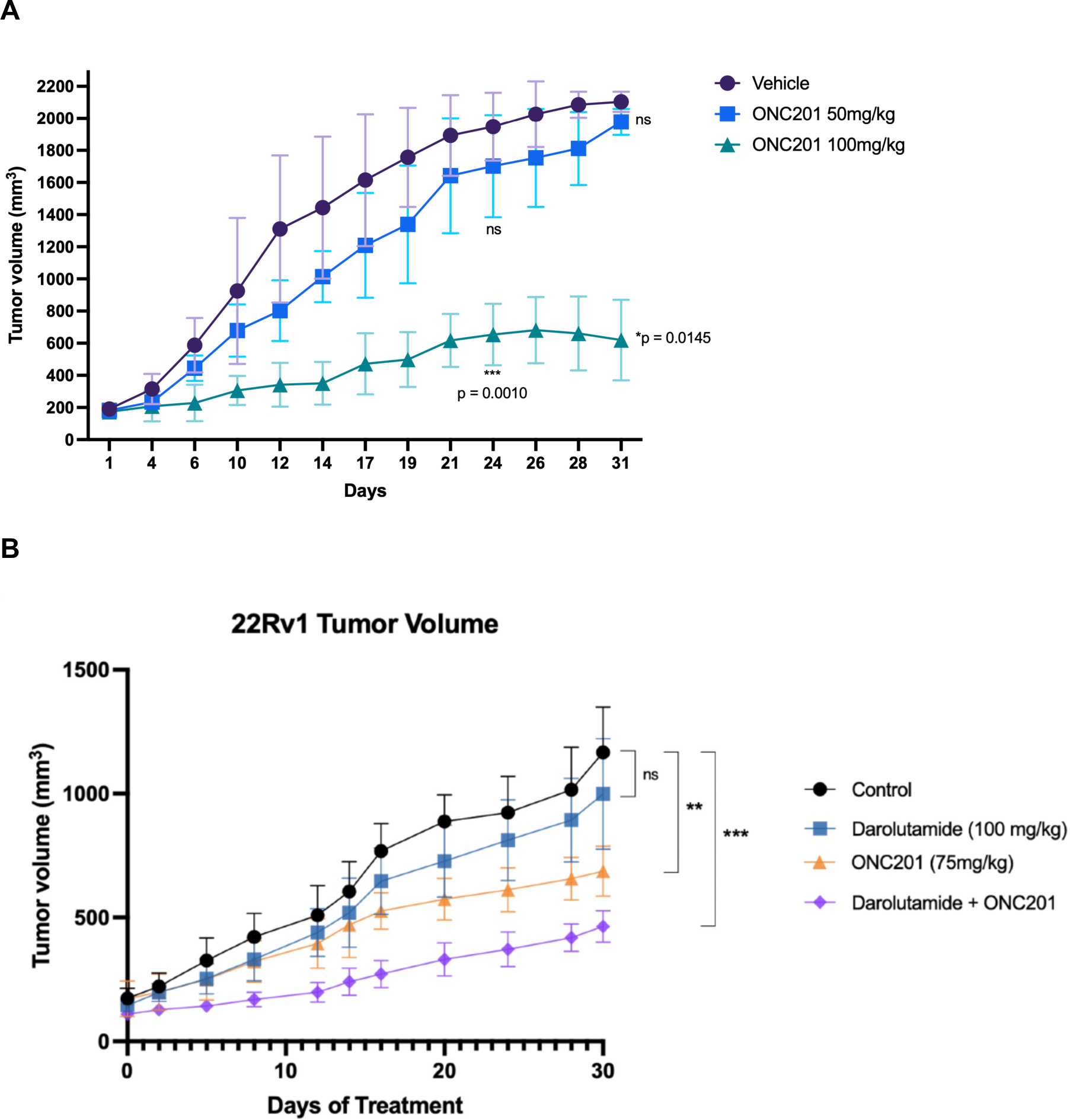
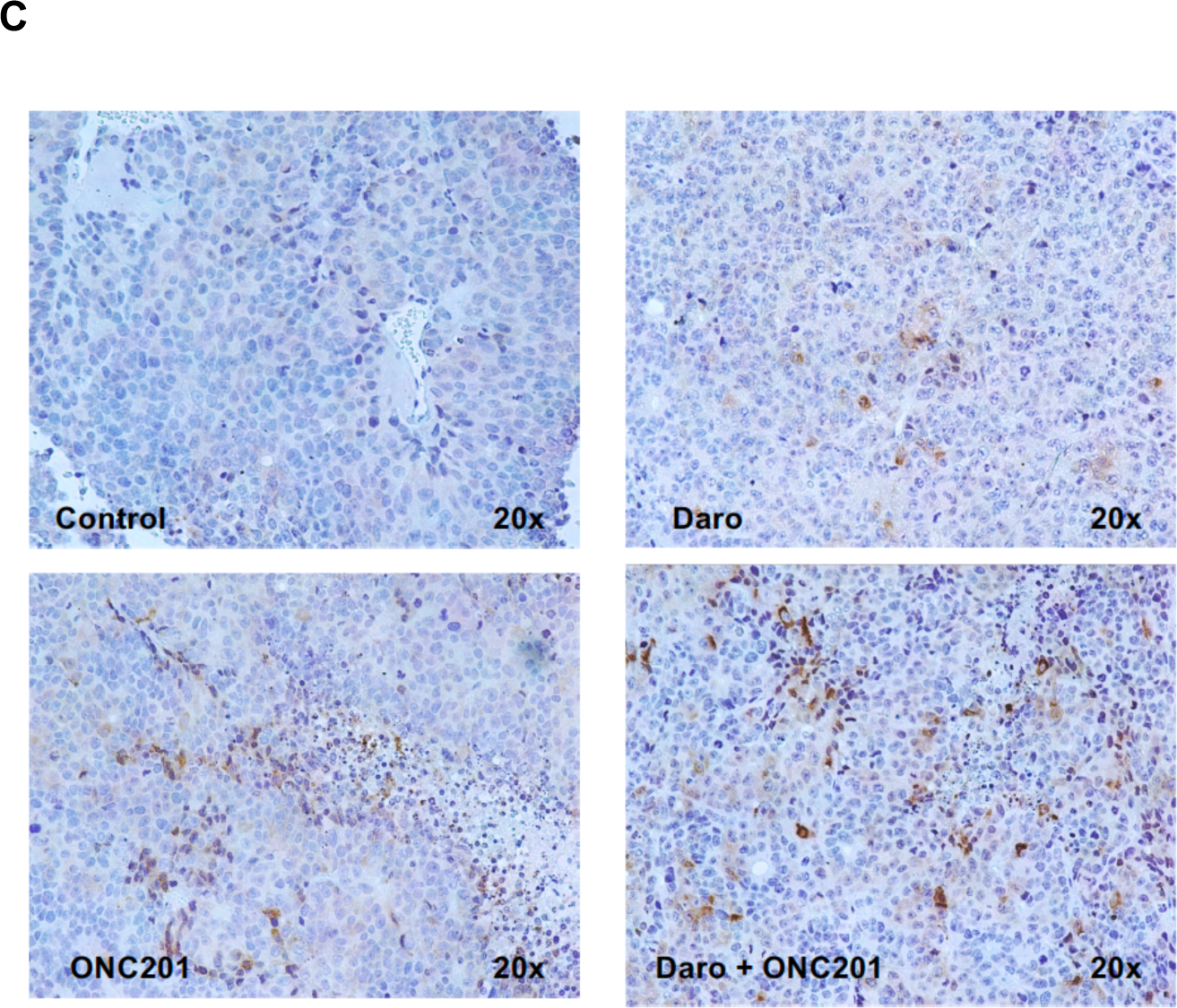
Growth of 22RV1 prostate cancer cells *in vivo*, anti-tumor efficacy of different doses of ONC201 alone or in combination with darolutamide along with increased TRAIL expression in tumor specimens. **(A)** Initial experiment with varying dose of ONC201 *in vivo* to treat 22RV1 xenografts. **(B)** Tumor growth delay of 22RV1 xenografts in vivo in control mice, ONC201- treated mice (75 mg/kg), darolutamide-treated mice (100 mg/kg) or the combination of ONC201 plus darolutamide. **(C)** Immunohistochemical stain for TRAIL protein expression in tumor xenografts from panel C under the different treatment conditions.

## Discussion

Our results demonstrate that a novel combination therapy using ONC201 and darolutamide is efficacious against human prostate cancer cells in culture and the combination shows therapeutic benefits in the 22RV1 *in vivo* xenograft model. The combination of ONC201 and both second generation NSAAs, enzalutamide and darolutamide, also displayed anti-proliferative and pro-apoptotic effects in prostate cancer cells regardless of AR expression and castration sensitivity. We show that both nmCRPC and mCRPC cell lines are sensitive to darolutamide and that darolutamide alone decreases PSA level, induces ATF4, and downregulates AR activity in prostate cancer cell lines regardless of hormone sensitivity, AR status, or the presence of AR mutations. We show that ONC201 reduces AR activity likely through the integrated stress response and subsequently reduces PSA level as single agent in prostate cancer cells. While both ONC201 and darolutamide show ATF4 upregulation 48 hours after treatment in the LNCaP cell line, significant reduction of PSA level is only observed in the combination groups. Since this observation was in the absence of PARP cleavage, hence no apoptotic activity, it was hypothesized that the induction of the ISR by the combination treatment and the competitive binding of darolutamide to the LBD of the AR collectively led to significant reduction in PSA level. We tested this hypothesis by co-treating the PCa cells with 1µg/mL DHT and the combination as well as single agent groups. Significant inhibition of AR activity was similarly observed in both LNCaP and 22RV1 cell lines treated with the darolutamide and ONC201 combination.

Initial analysis of therapeutic benefits of combining ONC201 at 50 mg/kg on a weekly schedule and the anti-androgens in prostate cancer xenograft models *in vivo* showed no statistical significance although the combination of ONC201 and darolutamide showed trends of partial antitumor growth arrest compared to the control group, especially after week 5 and towards the end of the six-week treatment period, demonstrated by reductions of final average tumor size (data not shown). Given that in our previous study where ONC201 administered three times a week with a dose of 50 mg/kg led to reduction in overall tumor sizes in both DU145 and PC3 xenograft models, the lack of robust anti-tumor effects with weekly ONC201 may be attributed to less frequent administration of the agent. We therefore tested 50 and 100 mg/kg ONC201 *in vivo* with 3x per week administration against 22RV1 xenografts and observed potent monotherapy effects with the 100 mg/kg dose. We therefore lowered the ONC201 dose to 75 mg/kg and observed in vivo synergy in combination with darolutamide.

Flow cytometry data showed trends of increased CD11b+ cells in treatment groups and TRAIL induction within NK cells after one week of treatment. The CD27 receptor is known to mediate the secretion of IFN-y [35, 36], yielding the highest IFN-y production in mature NK cells. Therefore, the trend of TRAIL upregulation seen could be induced by migrated intratumor NK cells, demonstrating early drug-induced immunomodulation in the tumor microenvironment. Notably, a clear upregulation of the activating receptor NKp46 on the cell surface of peripheral blood NK cells (or homing NK cells) was observed in the ONC201 and darolutamide combination group.

Although largely heterogenous, cytokine profiling results showed overall lower serum concentrations of tumorigenic cytokines IGF-1 and BAFF/Blys/TNFSF13B, and higher concentrations of immuno-modulatory cytokines such as IL-16, MMP8, TWEAK/TNFSF12 and Dkk-1. We then further divided the groups into partial responders to treatments and non-responders to investigate the potential roles of NK cells in solid tumors. Statistically significant higher concentrations of immune-modulatory cytokines IFN-y, GM-CSF, MMP-12, IL- 1 alpha, IL-2, and IL-3 were observed in the partial responders. However, little is known concerning the NK cell infiltration in solid tumors. Additionally, due to the possession of several subsets, NK cells have shown both immuno-activating and immuno-tolerating behaviors that are dependent on the specific organ types and receptors in specialized conditions [37, 38]. Further analyses are required to investigate the potential NK cell anti-tumor activities in the future *in vitro* and *in vivo*.

In summary, our present studies document a novel synergistic combination using ONC201 or ONC206 and darolutamide in treatment for prostate cancer. Cell culture data demonstrate significant drug sensitivity and synergistic anti-tumor efficacies in both metastatic and non-metastatic prostate cancer cell lines regardless of castration sensitivity and AR status. Synergistic effects are observed in AR negative cell lines, such as PC3 and DU145. *In vivo* experiments using 22RV1 CRPC cells with an optimized treatment and dosing regimen demonstrate therapeutic benefits of the combination therapy and roles of immune modulations against advanced prostate cancer. Our results warrant further translational and clinical studies with imipridones ONC201 or ONC201 in combination with enzalutamide or darolutamide for treatment of castrate resistant advanced or metastatic prostate cancer.

## Acknowledgements

W.S.E-D. is an American Cancer Society Research Professor and is supported by the Mencoff Family University Professorship at Brown University. This work was supported by an NIH grant (CA173453) to W.S.E-D. This work was presented in part at the 2022 meeting of the American Association for Cancer Research.

